# Ancestral Regulatory Mechanisms Specify Conserved Midbrain Circuitry in Arthropods and Vertebrates

**DOI:** 10.1101/820555

**Authors:** Jessika C. Bridi, Zoe N. Ludlow, Benjamin Kottler, Beate Hartmann, Lies Vanden Broeck, Jonah Dearlove, Markus Göker, Nicholas J. Strausfeld, Patrick Callaerts, Frank Hirth

## Abstract

Corresponding attributes of neural development and function suggest arthropod and vertebrate brains may have an evolutionarily conserved organization. However, the underlying mechanisms have remained elusive. Here we identify a gene regulatory and character identity network defining the deutocerebral-tritocerebral boundary (DTB) in *Drosophila*. We show this network comprises genes homologous to those directing midbrain-hindbrain boundary (MHB) formation in vertebrates and their closest chordate relatives. Genetic tracing reveals that the embryonic DTB gives rise to adult midbrain circuits that in flies control auditory and vestibular information processing and motor coordination, as do MHB-derived circuits in vertebrates. DTB-specific gene expression and function is directed by cis-regulatory elements (CREs) of developmental control genes that include homologs of mammalian *Zinc finger of the cerebellum* and *Purkinje cell protein 4*. Moreover, *Drosophila* DTB-specific CREs correspond to regulatory sequences of human *ENGRAILED-2, PAX-2* and *DACHSHUND-1* that direct MHB-specific expression in the embryonic mouse brain. Together, these findings imply ancestral regulatory mechanisms mediating the genetic specification of midbrain-cerebellar circuitry for balance and motor control that may predated the radiation of cephalic nervous systems across the animal kingdom.

## INTRODUCTION

Many components of the vertebrate and arthropod forebrain correspond with regard to their neural arrangements along the rostrocaudal axis and to the connections of higher integrative centres with sensory and motor pathways. In the ancient lineages from which derive, lampreys and hagfish, both proxies for ancestral vertebrates, the rostral neuropils of the forebrain encode visual and olfactory information relayed to further forebrain centers that integrate these modalities (1,2). The same arrangements apply to crown arthropods, amongst which Onychophora offer comparable proxies of ancestral neural arrangements (3). In contrast, circuits involved in vestibular reception and integration, and by extension acoustic perception, are located in more caudal territories of the anterior brain that arise in the telencephalon of vertebrates and in the deutocerebrum of arthropods (4,5).

It has been postulated that comparable brain organisation in arthropods and vertebrates is an example of genealogical correspondence, traceable to a distant pre-Cambrian ancestor by virtue of the conserved action of suites of homologous developmental control genes along anterior-posterior (AP) and dorso-ventral (DV) axes of the embryonic nervous system (6-13). For example, the *Drosophila* gene *orthodenticle (otd)* and its mammalian *Otx* homologues are required, respectively in the fly and mouse, for rostral brain development (14,15). In cross-phylum rescue experiments, human *OTX2* restores fly brain formation in *otd* mutant embryos (16,17) while fly *otd* can replace *Otx1/2* in mouse head and forebrain formation (18,19). Fly *engrailed* can replace *Engrailed-1* in mouse midbrain-hindbrain boundary (MHB) development (20). Cross-phyletic studies further revealed corresponding patterns of developmental genetic mechanisms, information processing and pathologies of the vertebrate basal ganglia and the arthropod central complex (9,21-23). These similitudes extend to comparisons of the vertebrate hippocampus and arthropod mushroom bodies, centers that support spatial navigation, allocentric memory, and associative learning (10,24).

Evidence from soft tissue preservation in fossils of stem arthropods, suggests that gross cerebral arrangements typifying the four extant panarthropod lineages originated latest in the early Cambrian, implying that neural ground patterns attributed to the Panarthropoda taxon may be both ancient and extremely stable over geological time (25). Here, ground patterns refer to ancestral arrangements that are inherited with modification. However, in the absence of detailed fossil material, resolving correspondences across phyla has to instead rely on the identification of shared developmental rules and their outcomes (21,24). These correspondences are expected to be defined by common gene regulatory (26) and character identity networks (27) that convey positional information and evidence of phenotypic homology, albeit often highly derived (28). Accordingly, it is cell identities, tissues and organs that yield information about common origins and identify corresponding ground patterns across lineages (29).

We applied this approach using developmental genetics to compare the formation and function of the *Drosophila* and vertebrate midbrain hindbrain boundary region. The vertebrate Midbrain-Hindbrain Boundary (MHB) is positioned by adjacent *Otx* and *Gbx* activity along the AP axis, and elaborated by region-specific expression of *Engrailed, Wnt, Pax2/5/8*, and FGF8-mediated organizer activity (30-33). In *Drosophila*, the corresponding boundary (henceforth referred to as the deutocerebral-tritocerebral boundary, DTB) is defined by comparable adjoining expression of *otd* and *unplugged (unpg)*, homologs of *Otx* and *Gbx*, respectively (34). The observation of these similar expression patterns raises a number of questions: whether they reflect a shared developmental program for the MHB and DTB; what adult brain structures derive from them; and what their function might be. We hypothesized that if the DTB evolutionarily corresponds to the vertebrate MHB, its formation would be mediated by gene regulatory and character identity networks homologous to those driving MHB formation. Furthermore, if true, we expected the DTB to provide circuits mediating behaviours commensurate with those regulated, and constrained, by MHB-derived circuitry. Here we describe experimental evidence verifying that the arthropod DTB indeed shares a ground pattern organization with the vertebrate MHB, including correspondence of neural circuits and their behavioral functions.

## RESULTS

We focused on phylotypic (35) stage 11-14 embryos to characterize morphological and molecular signatures of the developing *Drosophila* DTB. In addition to the adjoining *otd* and *unpg* expression and function reported earlier (34), we found specific domains of expression of the Pax2/5/8 homologs *shaven(sv)/dPax2* and *Pox neuro* (*Poxn*), as well as *engrailed* (*en*), *wingless* (*wg/dWnt*), *muscle specific homeobox* (*msh/dMsx*), *ventral nervous system defective* (*vnd/dNkx2*), and *empty spiracles* (*ems/dEmx*) (**Fig. 1A-C** and ***SI Appendix* Fig. S1A**). For axial patterning, we examined the expression and function of the key genes *otd* + *wg* (antero-posterior) and *msh* + *vnd* (dorsal-ventral), which revealed essential roles in DTB formation (***SI Appendix* Fig. S1B**).

**Figure 1.**
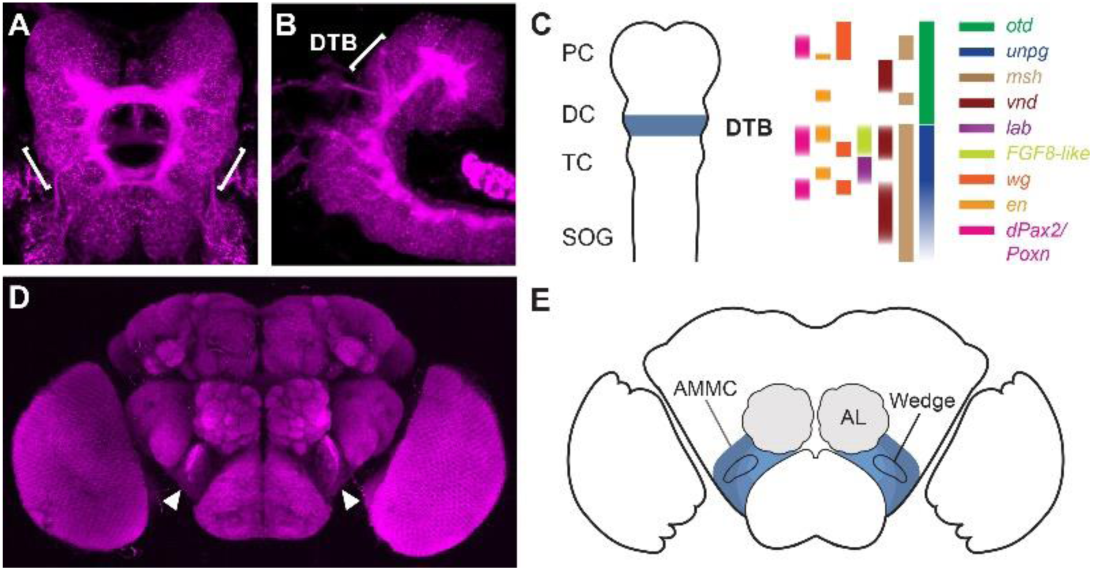
The embryonic deutocerebral-tritocerebral boundary gives rise to the antennal mechanosensory motor centre in the adult brain of *Drosophila*. (A, B, D) Confocal images of stage 14 embryonic (A, dorsal; B, lateral) and adult brain (D, frontal) immunolabeled with anti-Brp/nc82; bracket indicates deutocerebral-tritocerebral boundary (DTB) region; arrowheads indicate antennal mechanosensory motor centre (AMMC). (C) Schematic summarizing gene expression patterns delineating the DTB in the embryonic brain; PC, protocerebrum; DC, deutocerebrum; TC, tritocerebrum; SOG, subesophaegal ganglion. (E), schematic of adult brain showing AMMC, Wedge and antennal lobes (AL).

A cardinal feature of the vertebrate MHB is its organizer activity, mainly mediated by the FGF8 effector molecule (30-33,36). Previous studies failed to identify a DTB-related function of the FGF8 homolog *branchless* and its receptor *breathless* in embryonic brain development of *Drosophila* (34). We now show that a second set of FGF8-like orthologs, *thisbe/FGF8-like1* and *pyramus/FGF8-like2* (37), and the FGF8 receptor heartless (*htl*), are expressed at the DTB (***SI Appendix* Fig. S2**). A functional role for FGF8 signalling at the DTB was revealed by altered *engrailed* expression patterns (**Fig. 2A**, arrows) and morphological defects affecting longitudinal connectives (**Fig. 2A**, arrowheads) in embryos homozygous for a deficiency, *Df(2R)BSC25*, uncovering both *FGF8-like1* and *FGF8-like2* genomic loci, and of a *htl* null allele. These observations were further substantiated by progressive changes and loss of the DTB expression patterns of *unpg* and *ems* in *htl* null mutant embryos, between embryonic stages 12-16 (**Fig. 2B**). These data identify a role of FGF8-like signalling in the maintenance of the embryonic DTB region in *Drosophila*. In contrast, ectopic expression of the FGF8-like homologue *thisbe* in *ems*-specific brain regions did not cause any detectable changes in morphology or molecular signatures of the DTB region (***SI Appendix* Fig. S3**). Despite conserved regulatory interactions between *otd/Otx* and *unpg/Gbx* (34), these data indicate the absence of FGF8-mediated organizer activity in the embryonic DTB. We conclude that FGF8-like signalling is required for the maintenance of the embryonic DTB, but contrary to what is seen in vertebrate MHB development appears not to organize the DTB region in *Drosophila*.

**Figure 2.**
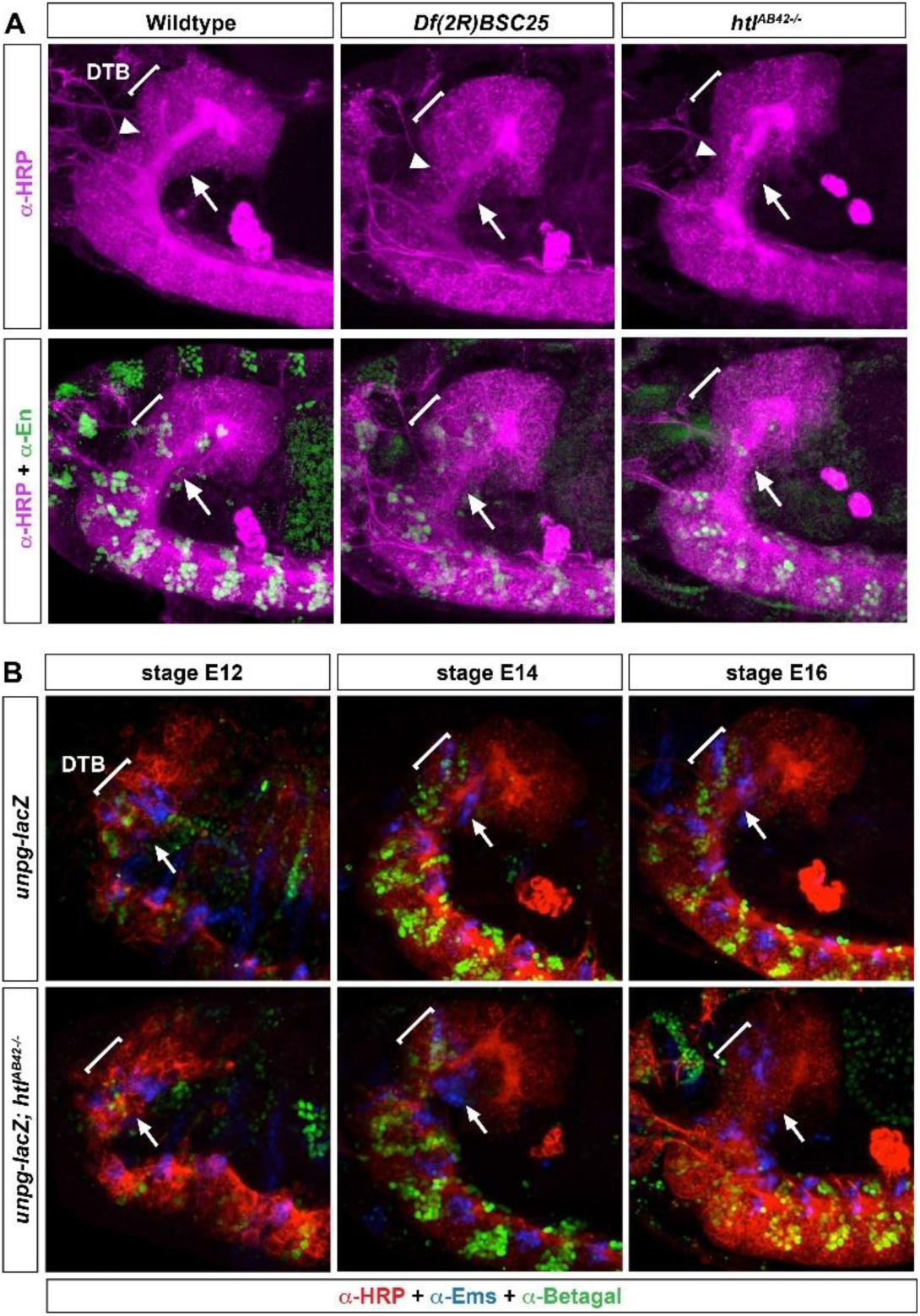
FGF8 signalling is required for the formation and maintenance of the embryonic DTB. Confocal images of stage 13/14 embryonic brains, in (A) immunolabelled for HRP (red) and anti-Engrailed (green/yellow), in (B) immunolabelled for HRP (red), anti-Ems (blue) and anti-β-galactosidase (green/yellow), lateral views; brackets and arrows indicate deutocerebral-tritocerebral boundary (DTB). (A) in wildtype, the DTB is characterized by a stripe-like engrailed expression pattern, that is affected in *Df(2R)BSC25*, a deficiency deleting both *FGF8-like1* and *FGF8-like2* genomic loci, as well as in *htl* null mutant. Note morphological defects, especially for longitudinal connectives of deutocerebral-tritocerebral neuromeres (arrowhead) (B), transgenic *unpg-lacZ* control brains at embryonic stages E12, E14 and E16, respectively. Progressive *unpg-lacZ* and *ems* expression patterns reveal successive formation of the DTB. However, in *htl* null mutant embryos (*unpg-lacZ; htl*^*AB42-/-*^), *unpg* and *ems* expression are initially visible but subsequently lost by embryonic stage 16).

To identify the adult brain structures and functional modalities that arise from the embryonic DTB, we used genetic tracing of neural lineages (38,39). We traced *engrailed* expressing lineages of the embryonic neuroectodermal DTB (***SI Appendix* Fig. S4A-C**) and identified neurons and projections of the antennal mechanosensory and motor centre (AMMC) and select antennal glomeruli in the adult brain (**Fig. 1D, E** and ***SI Appendix* Fig. S4D-F**). We also determined the fate of Poxn expressing cells which in the embryonic brain are located next to DTB-specific engrailed lineages (**Fig. 3A-H**), and in the Wedge of the adult brain where, similar to En+ cells, they express the neurotransmitter GABA (**Fig. 3I-P**). Genetic tracing revealed that Poxn-expressing AMMC/Wedge neurons derive from DTB lineages, with their region-specific projection patterns resembling the mechanosensory pathway architecture of local interneurons and projection neurons (**Fig. 3Q-T**) previously identified for the AMMC and the Wedge (40-43).

**Figure 3.**
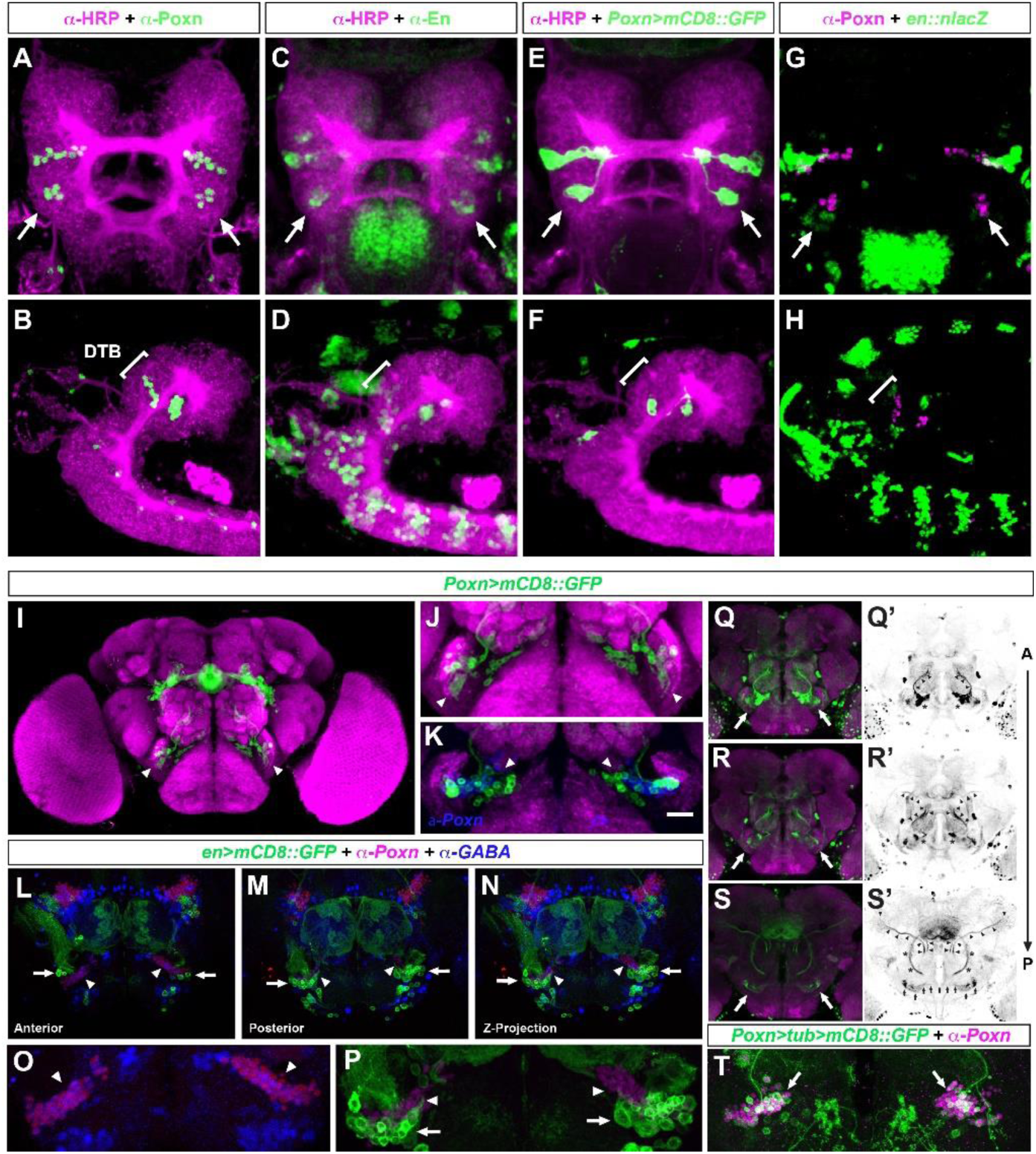
DTB and AMMC/Wedge-specific expression of the Pax2/5/8 homologue *Pox neuro*. Confocal images of embryonic stage 14 (A-H); anterior is up in A, C, E, G, dorsal views; anterior is to the left in B, D, F, H, lateral views. (A, B), in the anterior embryonic brain, anti-Pox neuro (Poxn) immunolabelling reveals two Poxn expression domains, an anterior at the protocerebral-deutocerebral neuromere boundary and a posterior demarcating the deutocerebral-tritocerebral boundary (DTB) region (arrows, bracket). (C, D), engrailed expression demarcates neuromere boundaries, including the DTB (arrows, bracket). (E, F) *Poxn>mCD8::GFP* expression reveals GFP expression pattern comparable to endogenous Poxn expression (compare with A, B), including the DTB (arrows; bracket). (G, H) *en>nLacZ* expression reveals Engrailed expression pattern comparable to endogenous En expression (compare with C, D), including the DTB that encompasses Poxn expression domain (arrows; bracket). (I-T), confocal images of adult brain; dorsal is up. (I-K) *Poxn*^*brain*^*>mCD8::GFP* mediated cell labelling identifies Poxn+ cell clusters (green, arrowheads) in close vicinity to the antennal mechanosensory motor centre (AMMC), the majority of which are anti-Poxn immuno-positive (K, in blue). (L-P) *en>mCD8::GFP* visualises AMMC neurons (arrows) that are located in close vicinity/adjacent to Poxn positive cells (magenta, arrowheads; enlarged views in O, P) that are immunoreactive for anti-GABA (O, blue, arrowheads) like En-expressing cells (P, arrows). (Q-S’), *Poxn*^*brain*^*>mCD8::GFP* visualises AMMC/Wedge neurons (arrows) and their projections to antennal glomeruli (Q, Q’) to ventrolateral protocerebrum (R-S’, middle section of brain), as well as commissural axons of AMMC/Wedge neurons (S’, small arrows). (T) *Poxn>tub>mCD8::GFP* mediated genetic tracing of DTB Poxn lineages identifies AMMC/Wedge neurons (arrows). Scale bar in K, 20μm.

The *Drosophila* AMMC and Wedge neuropils comprise neurons that mediate auditory, vestibular, mechanosensory and somatosensory information processing in pathways with similarities to the mammalian auditory and vestibular pathways (40-43). In vertebrates, auditory, vestibular, somatosensory and motor information, are processed by neural populations of the tectum and cerebellum, adult brain structures derived from the MHB region (36, 44). The tectum and cerebellum receive auditory and vestibular, as well as motor information and, among other functions, are important for balance, body posture, sensorimotor integration and motor coordination (1,4,45).

To test whether DTB-derived circuits in *Drosophila* might exert similar functions, the GAL4-UAS system was used to express *Tetanus-Toxin-Light-Chain* (*TNT*) and inhibit synaptic transmission (46) in subsets of AMMC neurons (41). Flies were tested for their startle-induced negative geotaxis (SING) response, which after being shaken to the bottom in a test tube quantifies their ability to right themselves and climb upwards (47). Except *R19E09>TNT*, all of the tested genotypes showed significantly impaired SING behaviour (**Fig. 4A** and ***SI Appendix* Table S1**), including R79D08, R45D07 and R30A07 that target AMMC neurons and co-express *engrailed*, or encompass *Poxn* and *engrailed* expression domains (**Fig. 4C-I**). Of note, several of the tested genotypes showed difficulties with balance and to right themselves, as exemplified for *R52F05>TNT* compared to control (***SI Appendix* Movies S1** and **S2**).

**Figure 4.**
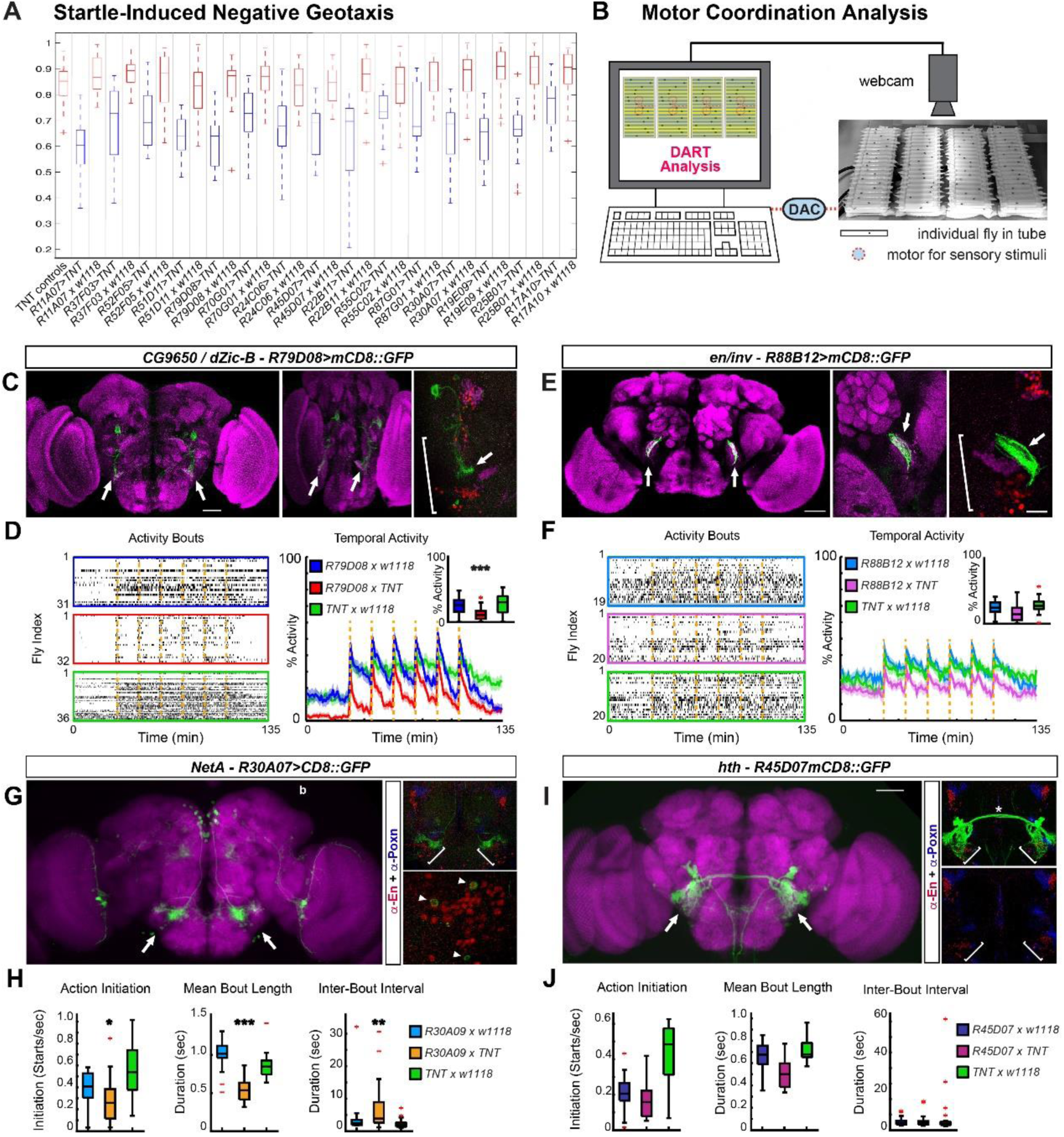
The AMMC mediates motor coordination in *Drosophila*. (A) Startle-induced negative geotaxis of AMMC-specific GAL4 lines misexpressing *UAS*-*TNT* and controls (n=150 each). (B) *Drosophila* Arousal Tracking (DART) recording each fly in tube walking back and forth, motors underneath elicit vibration stimuli via digital-to-analogue converter (DAC). (C) *CG9560/dZic-B, R79D08>mCD8::GFP* immunolabeled with anti-Brp/nc82 and anti-GFP visualizes AMMC-specific (bracket) arborizations (arrows); middle panel, rotated brain to depict AMMC-specific projections. (D) Motor behaviour of *R79D08>TNT*, UAS/+ and Gal4/+ control flies; left, raster plots of activity bouts, each lane one individual fly, coloured boxes indicate genotypes; right, stimulus response (main plot) and median response (inset) to repeated mechanical stimulation (dashed orange lines). (E) *inv R88B12>mCD8::GFP* visualizes neuronal projections to AMMC (arrows, enlarged for hemisphere in middle and right panel) encompassed by anti-En (green) and anti-Poxn (magenta) expression domains (bracket). (F) Motor behaviour of *R88B12>TNT*, UAS/+ and Gal4/+ control flies, parameters as in D. (G) *NetA, R30A07>mCD8::GFP* expression in AMMC (left, arrows); right, anti-En (red) and anti-Poxn (blue) immunolabelling encompasses *R30A07>mCD8::GFP* domain (brackets) and cells, some co-expressing Engrailed (arrowheads). (H) Motor kinematics of *R30A07>TNT*, UAS/+ and Gal4/+ control flies. (I) *hth R45D07*>m*CD8::GFP* in AMMC (arrows); right, *R54D07>mCD8::GFP* visualises AMMC interneurons and dendritic arborisations close to anti-Engrailed (red) and anti-Poxn (blue) immunolabelled neurons (blue) that encompass AMMC/Wedge (brackets). (J) Motor kinematics of *R45D07>TNT*, UAS/+ and Gal4/+ control flies. Mean ± Standard Error of the Mean (SEM), asterisks, p<0.05 (*), p<0.01 (**) or p<0.001 (***). Scale bars, 50μm (C, E, left) and 10μm (E, right).

To further analyze AMMC-mediated motor coordination, we employed video-assisted motion tracking and recorded freely moving flies (**Fig. 4B**). During 135-minute recordings, activity bouts and movement trajectories were analyzed to quantify locomotion parameters: frequency of episodic movements, how often they were initiated, their length and average velocity, as well as the duration and frequency of pauses. Response to sensory stimulation triggered by repeated mechanical shock were also recorded. *UAS-TNT* expression by *R79D08* (*CG9650/dZic-B*)-Gal4, which targets zone B (40-42) of the AMMC (**Fig. 4C**, arrows), significantly impaired overall activity and duration, with fewer actions initiated, shorter episodes of activity and their intervals, reduced velocity and distances travelled (**Fig. 4D** and ***SI Appendix* Fig. S5A**). *UAS-TNT* expression by *R88B12* (*en/inv*)-Gal4 targeting zone A (40-42) of the AMMC (**Fig. 4E**, arrows) significantly impaired average and pre-stimulus speed, with shorter bouts of activity, together resulting in less distance travelled (**Fig. 4F** and ***SI Appendix* Fig. S5B**). Comparable motor phenotypes were seen with *R30A07* (*NetA*)-Gal4, which targets AMMC neurons that co-express *engrailed* (**Fig. 4G**, arrowheads; **Fig. 4H** and **Fig. S5C**), but not with *R45D07* (*hth*) Gal4 targeting parts of the AMMC-specific giant fiber system (**Fig. 4I, J** and ***SI Appendix* Fig. S5D**). Together, with SING data, our behavioural observations establish essential functions of the AMMC for sensorimotor integration (40,41,43), balance, righting reflex and motor coordination in *Drosophila*, behavioural manifestations similar to MHB region-derived circuits and their ancestral functions.

Our findings thus far establish correspondences between *Drosophila* DTB and vertebrate MHB at multiple levels including adult brain circuits and the behaviours they regulate. We hypothesized that this will be reflected in commonalities among character identity networks of DTB and MHB that are mediated by homologous gene regulatory networks (27,28). To test this hypothesis, we screened the Janelia Gal4 collection (48) for cis-regulatory elements (CREs) mediating the spatio-temporal expression of developmental genes controlling DTB formation in *Drosophila*. We identified CREs for *msh, vnd, ems*, and for *thisbe/FGF8-like1* (***SI Appendix* Fig. S6A-E)**, genes that are essential for the formation and/or maintenance of the embryonic DTB (**Fig. 2** and ***SI Appendix* Fig. S1** and ref. 14). In addition, we identified CREs for *Wnt10, Sex combs reduced/Hox5;* the *Drosophila* homologs of *zinc finger of the cerebellum (ZIC), odd-paired* (*opa/dZic-A*) and *CG9650/dZic-B;* of *Purkinje cell protein 4* (*PCP4*), *igloo* (*igl/dPCP4*); of *Ptf1a, Fer1/dPtf1a*, and of *Lim1* (***SI Appendix* Fig. S6F-J**). All mammalian homologs of these genes have been implicated in vertebrate MHB formation and the specification of midbrain-cerebellar circuitry (30-34, 49) as shown in ***SI Appendix* Table S2**.

We also identified CREs for *dachshund* (*dac*) and the *Pax2* homologue *shaven* (*sv/dPax2*). Consistent with *engrailed/invected* and *Poxn*-related genetic tracing of DTB-AMMC lineages, *dac*-specific CRE *R65A11-Gal4* targeted *UAS-mCD8::GFP* expression to the procephalic DTB region, in derived lineages of the embryonic brain and to the AMMC in a pattern encompassed by DTB-specific *engrailed* and *Poxn* expression domains (**Fig. 5A-C**). The regulatory element VT51937 (50) located within an intronic region of the *sv/dPax2* locus targets Gal4 expression in a segment-specific pattern similar to endogenous *sv/dPax2*, including DTB expression domains (***SI Appendix* Fig. S7A-F**). *VT51937-Gal4* mediated genetic tracing also identified cells and projections in the AMMC (***SI Appendix* Fig. S7G**). Together these data identify CREs of the DTB-AMMC character identity network in *Drosophila* that mediate the spatiotemporal expression patterns of genes that are homologous to genes involved in the formation and specification of the vertebrate MHB and derived midbrain-cerebellar circuitry (30-34,49).

**Figure 5.**
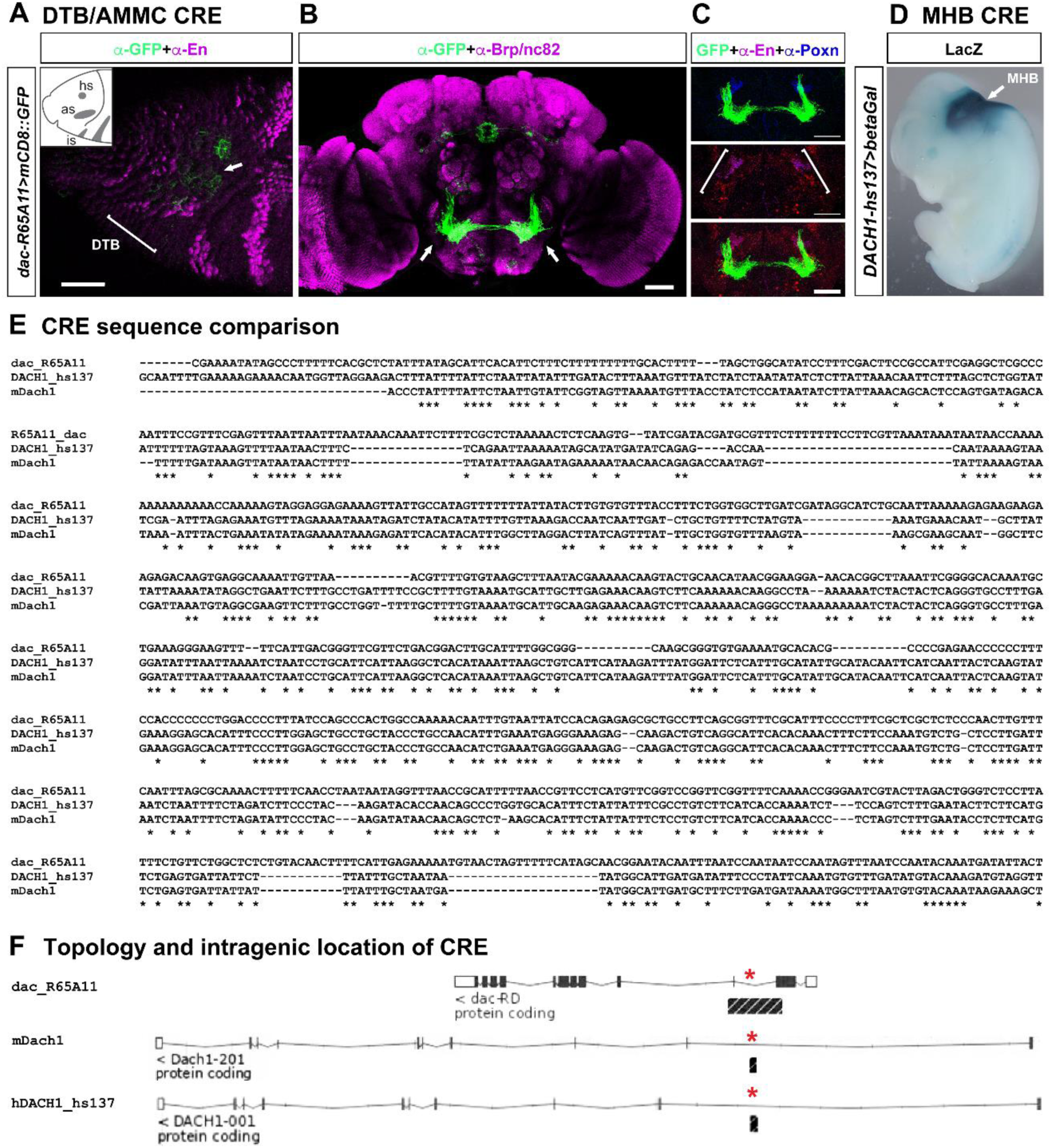
Conserved cis-regulatory sequences of *dac/DACH1* direct DTB-AMMC specific expression in *Drosophila* and MHB-specific expression in mouse. (A) Confocal image of stage 10/11 *Drosophila* neuroectoderm of *R65A11-Gal4>mCD8::GFP* embryo (lateral view, anterior left, dorsal up), immunolabeled with anti-GFP (green) and anti-En (magenta); inset illustrates Engrailed expression domains in procephalic neuroectoderm including head spot (hs), antennal spot (as) and intercalary spot (is). *R65A11>GFP* expression (arrow) is seen in DTB primordium (bracket). (B) Confocal image of *R65A11-LexA>mCD8::GFP* expression in AMMC (arrows) of adult *Drosophila* brain, immunolabeled with anti-Brp/NC82 (magenta) and anti-GFP (green). (C) Anti-En and anti-Poxn immunolabeling encompass *R65A11-LexA>mCD8::GFP* in AMMC (bracket). (D) human *DACH1*-specific cis-regulatory sequence (CRE), *hs137*, targets LacZ expression to midbrain hindbrain boundary (MHB, arrow) in E11.5 mouse embryo (VISTA database). (E) Sequence comparison of parts of *D. melanogaster dac R65A11*, mouse *mDach1* and human *hDACH1 hs137* CREs. (F) Intragenic locations (black bar, asterisks) of *dac R65A11, hDACH1 hs137* and corresponding mouse *mDach1* CRE sequence.

Comparable to the observed genealogy of DTB-AMMC lineages in *Drosophila*, studies on murine brain development showed that tectum, tegmentum and cerebellar Purkinje and granule cells in the adult brain derive from the embryonic MHB (36,44). Given the correspondences between the *Drosophila* DTB and vertebrate MHB gene regulatory and character identity networks, we asked whether cis-regulatory elements are conserved. To determine potential cross-phylum CRE conservation, we utilized DTB-AMMC specific regulatory sequences and applied bioinformatics tools (51) including VISTA (52), MLAGAN (53) and EMBOSS MATCHER (54) to screen for corresponding CREs in mouse and human genomes (55). Stringent selection criteria (56) were applied to identify CRE sequences that are *(i)* linked to the same homologous genes in the different species; *(ii)* there is a minimum of 62% sequence identity over at least 55 base pairs with minimum 1e^-1^ confidence level; *(iii)* the CREs are not un-annotated protein sequences; and *(iv)* the CREs are not repetitive elements.

We first analyzed the DTB-AMMC-specific CRE of *sv/dPax2* (= VT51937 sequence) and identified non-coding CREs for mouse *Pax2* and human *PAX2* with extensive sequence similarities (***SI Appendix* Fig. S7H** and **data set S1**), and comparable intragenic location (***SI Appendix* Fig. S7I**). Following this strategy, we used DTB-AMMC-specific CRE sequences for *dachshund* and *engrailed/invected* and identified corresponding human CREs conserved among vertebrates that direct MHB-specific expression in mouse for the *dachshund* homologs *Dach1/DACH1* (**Fig. 5D-F**) and for the *engrailed/invected* homologs *Engrailed2/EN2* (***SI Appendix* Fig. S8**). Further bioinformatics analysis identified core CRE sequences associated with *dac* and *DACH1, en/inv* and *EN2* and *sv* and *PAX2* in *Drosophilidae* and vertebrate genomes (***SI Appendix* data sets S1-S3**), suggesting ancestral non-coding regulatory sequences and their function predate the radiation of insect-specific DTB and vertebrate-specific MHB circuits and morphologies.

## DISCUSSION

We have identified gene regulatory and character-identity networks that underlie the formation of the deutocerebral-tritocerebral boundary in *Drosophila*. Mutant analyses reveal that *otd* + *wg* and *msh* + *vnd*, acting along the AP and DV body axes, respectively, are required for the formation of the embryonic DTB, and that FGF8-like signaling is necessary for its developmental maintenance. Genetic tracing experiments, together with the analysis of cis-regulatory elements for *engrailed/invected, dachshund* and *shaven/dPax2*, as well as behavioral analysis after synaptic inactivation show that embryonic DTB lineages give rise to neural circuits in the AMMC/Wedge complex of the adult brain that mediate balance and motor coordination in *Drosophila*. Together these findings establish a ground pattern of DTB formation and derived circuit function in *Drosophila* that corresponds to the ground pattern, gene regulatory and character identity networks of the vertebrate MHB and derived midbrain-cerebellar circuits (***SI Appendix* Fig. S9**).

These data imply that caudal domains of the arthropod deutocerebrum and its circuits in *Drosophila* correspond to the vertebrate MHB and its derived principle proprioceptive circuits (see ***SI Appendix* Fig. S9**). It must be emphasized here that these are not ascribed to the cerebellum, the anlage of which forms as an asegmental volume within Gbx2 and non-Hox expression domains of the developing MHB (30-33,36,57). In vertebrates, FGF8 signalling acts as a secondary organizer in boundary development of the MHB and by promoting growth essential for the formation of tectum and cerebellum (30-34,36, 57). We did not observe FGF8-like organizer activity in flies but a role in the maintenance of the DTB boundary, suggesting an ancestral boundary-related function for FGF8 (57). Yet the absence of extended proliferative activity in *Drosophila* (34 and this paper) suggests that growth-related organizer activity of FGF8 is a vertebrate innovation, whereas the boundary region is defined by expression patterns of genes homologous to those observed at the DTB/MHB (**Fig. 6**). Indeed, in spite of comparable expression domains (11,12,58, 59), no phenotypic cerebellum is found in ascidians, hemichordates and cephalochordates; none of which can be assumed as proxies for ancestral vertebrates, but all of which may as likely represent highly derived and evolutionary simplified crown species. However, ancestral circuits mediating vestibular and motor (balance) coordination, which are specified by genes and regulatory networks homologous to those described in this paper, can be found in the persisting ancient lineages of early vertebrates, lampreys and hagfish (2).

**Figure 6.**
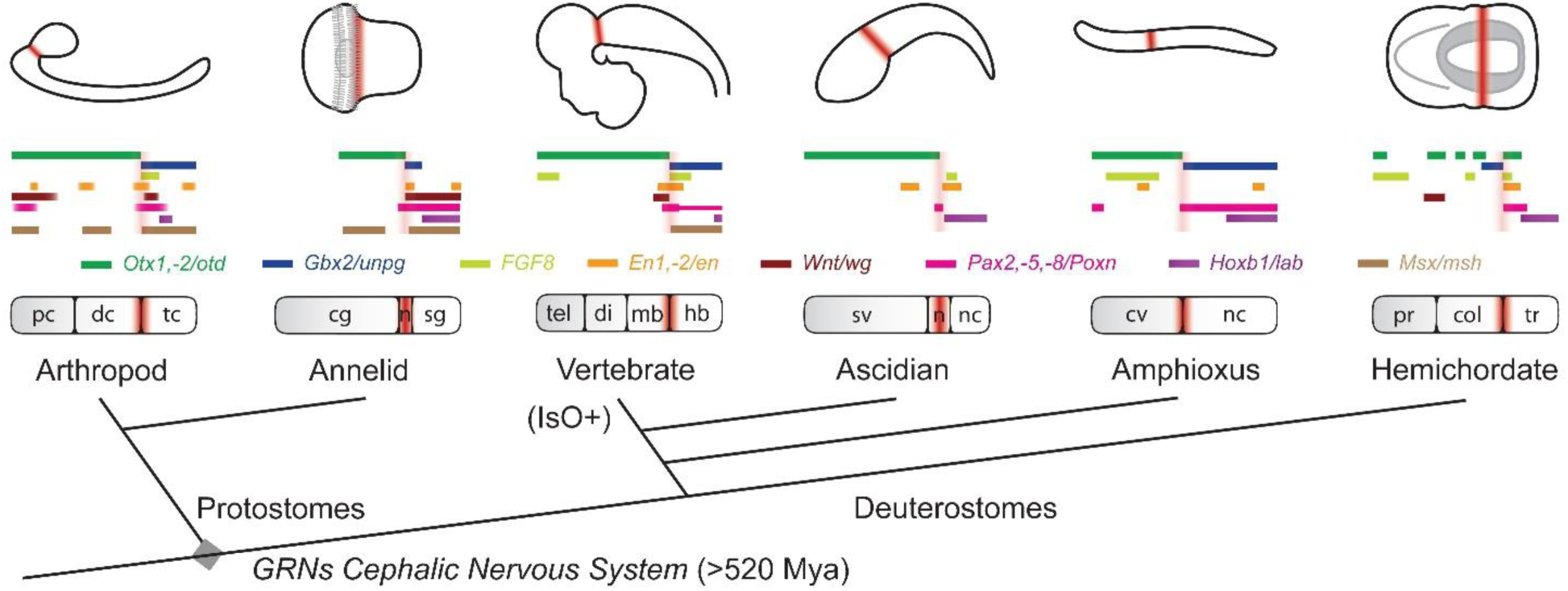
Phylogenetic comparison of DTB/MHB-related molecular signatures in nervous systems of extant Bilateria. Schematic diagram of homologous gene expression in central nervous system of arthropod *Drosophila melanogaster*, annelid *Platynereis dumerilii*, vertebrate *Mus musculus*, ascidian *Ciona intestinalis*, amphioxus *Branchiostoma floridae*, and ectodermal nervous system of hemichordate *Saccoglossus kowalevski*. Embryos and bar diagrams are oriented anterior to the left, dorsal up; brown colouring indicates boundary region. Multi-level correspondences of arthropod DTB and vertebrate MHB ground pattern organization suggest ancestral origin of gene regulatory networks (GRNs) for cephalic nervous systems >520 million years ago (Mya) that predate the radiation into protostomes and deuterostomes, and a suggested origin of the MHB-specific isthmic organizer (IsO) in the vertebrate lineage. Abbreviations: cg, cerebral ganglion; col, collar; dc, deutocerebrum; di, diencephalon; hb, hindbrain; mb, midbrain; Mes, Mesencephalon; Met, Metencephalon; Mye, Myelencephalon; n, neck; nc, nerve cord; pc, protocerebrum; pr, proboscis; sg, segmental ganglia; SOG, Subesophageal ganglion;sv, sensory vesicle; tc, tritocerebrum; tel, telencephalon; tr, trunk.

The observed correspondences in circuit formation extend to behaviours they regulate. UAS-TNT expression mediating synaptic inactivation, for example in *R88B12-Gal4* and *R30A07-Gal4* targeting DTB-derived sub-circuits of the AMMC/Wedge complex, results in flies with impaired balance, defective action initiation and maintenance, and compromised sequences of motor actions. These AMMC circuits have been shown to mediate auditory and vestibular information processing and coordination (40,41), suggesting the DTB-derived AMMC/Wedge circuits integrate mechanosensory submodalities for motor homeostasis. These functions correspond to activity of MHB-derived acoustic and vestibular receptor pathways in vertebrates (40,43) which, when impaired in inherited disorders, affects both auditory and vestibular functions such as seen in ataxic patients (60). The observed correspondences therefore suggest that similar to MHB-derived circuits and their ancestral functions, the DTB-derived AMMC/Wedge circuits in Arthropoda are required for sensorimotor integration, body posture and motor coordination. These findings identify correspondences between ground patterns of the insect DTB and vertebrate MHB that extend beyond homologous genes and their spatio-temporal expression patterns and functions, to neural circuits and behavior.

The present results identify CREs with extensive sequence similarities associated with highly conserved developmental control genes regulating boundary formation between the rostral brain and its genetically distinct caudal nervous system in insects and vertebrates. Core elements of the identified CREs are conserved and are employed for the formation of corresponding circuits and roles in neuronal processing. In conclusion, the corresponding ground patterns of insect DTB and vertebrate MHB suggest the early appearance in bilaterian evolution of a cephalic nervous system that evolved predictive motor homeostasis before the divergence of the protostome lineages and before the origin of deuterostomes. The observed correspondences therefore hypothesize the retention across phyla of conserved regulatory mechanisms (61-64) necessary and sufficient for the formation and function of neural networks for adaptive behaviors (22) common to all animals that possess a brain.

## MATERIALS AND METHODS

### Fly Strains and Genetics

The wild-type strain used was Oregon R. The following mutant alleles and characterization constructs were used to investigate expression and function: *P{en2*.*4-Gal4}e16E, UAS-mCD8::GFP*^*LL5*^ *and poxn*^*brain*^*-Gal4* as well as *UAS-mCD8::GFP, tub-FRT-CD2-FRT-Gal4, UAS-FLP/CyO GMR Dfd YFP* (65); *otd*^*JA101*^ (14); *P{lacZ}unpg*^*f85*^ (an *unpg-lacZ* reporter gene that expresses cytoplasmic β-galactosidase in the same pattern as endogenous *unpg*) (34); *P{lacZ}Pax2*^Δ*122*^ (a *Pax2-lacZ* reporter gene that expresses β-galactosidase in the same pattern as endogenous *Pax2*) (34); *P{3*′*lacZ}unpg*^*r37*^ (*unpg* null allele with a *unpg-lacZ* reporter gene that expresses nuclear β-galactosidase in the same pattern as endogenous *unpg*) (34); *wg*^*CX4*^ and *wg-lacZ* (Bloomington); *msh*^Δ*68*^ *(66); vnd*^*6*^ *(67);* the deficiency *Df (2R) BSC25* that removes the FGF8-like 1 and FGF8-like 2 loci together with adjacent regions and *htl*^*AB42*^ (37) *UAS-FGF8-like 1* (37); and *ems2*.*6 (72*.*5)-Gal4* (S.G. Sprecher, unpublished).

To generate *Poxn*^*brain*^*-Gal4* flies, the *Poxn* brain enhancer (68) was amplified by PCR from genomic DNA. The PCR product was subcloned into *pPTGal* vector using *XbaI* and *NotI* sites, followed by sequencing; the genomic region 2R:11723830 to 11725559 was inserted into *pPTGal*. Primer sequences are:

forward, *5’-gctcattaatgaccatgaaa-3’*;

reverse, *5’-aagcggccgcgttaagtaacgctcggtgg-3’*.

Transgenesis was performed by BestGene Inc (CA, USA).

For lineage tracing, the following strains were used: *w*^*1118*^ (control), *P{en2*.*4-Gal4}e16E, UAS-mCD8::GFP*^*LL5*^, or *poxn*^*brain*^*-Gal4* were crossed to *UAS-mCD8::GFP, tub-FRT-CD2-FRT-Gal4, UAS-FLP/CyO GMR Dfd YFP*. Offspring were raised at 18°C to suppress random leaky FLP activity.

For behavioural experiments, we used *UAS-TNT-E* (46) crossed to AMMC-specific *Gal4* lines. Corresponding controls for *Gal4* driver and for UAS responder line, were generated by backcrossing to *w*^*1118*^. All behavioural experiments were carried out in a temperature-controlled chamber at 25°C.

### In situ Hybridization, Immunocytochemistry and Image Analysis

For in situ hybridization experiments, digoxigenin-labelled sense and antisense RNA probes were generated in vitro with a DIG labelling kit (Roche diagnostics) and hybridized to *Drosophila* whole-mount embryos, following standard procedures (69).

Whole-mount immunocytochemistry was performed as previously described (70,71). Primary antibodies were rabbit anti-Otd (34), used 1:100; rabbit anti-Msh (66), used 1:500; rabbit anti-Vnd (67), used 1:200; rabbit anti-sv/dPax2 (34), used 1:50; monoclonal anti-Poxn antibodies (72), used 1:20; rabbit anti-HRP (FITC-conjugated, Jackson Immunoresearch), used 1:50; mouse anti-En (Developmental Studies Hybridoma Bank, DSHB), used 1:1; rabbit anti-β-Gal, used 1:200 (Milan analytica); mouse anti-β-Gal (DSHB), used 1:100; rabbit anti-Lab (73), used 1:50; rat anti-Ems (74), used 1:2000; mouse anti-Bruchpilot (nc82, DSHB), used 1:20; mouse anti-Synapsin (3C11, DSHB), used 1:50; rabbit anti-GFP (Invitrogen), used 1:500; rabbit anti-GABA (Sigma-Aldrich), used 1:1000. Secondary antibodies were Alexa-568-conjugated goat anti-mouse, Alexa-568-conjugated goat anti-rabbit, Alexa-568-conjugated goat anti-rat, Alexa-488-conjugated goat anti-mouse, Alexa-488-conjugated goat anti-rabbit, and Alexa-488-conjugated goat anti-rat (Molecular probes), all used 1:150. Embryos, larval CNS and adult brains were mounted in Vectashield H-1000 (Vector).

Fluorescence samples were scanned and recorded with a Leica TCS SP5 confocal microscope. Z-projections were created and analysed using FIJI. Images were processed using Adobe Photoshop and figures constructed in Adobe Illustrator.

### Startle Induced Negative Geotaxis Assay (SING)

Groups of 10 flies with shortened wings of the same age, sex and genotype were placed in a vertical column (19 cm long, 2cm diameter). The wings were clipped under sedation (with CO2) at least 24 hours prior to testing. They were suddenly startled by gently tapping them down, to which *Drosophila* responds by climbing up. After 10 seconds, it was counted how many flies were above the 2cm mark and the trial was repeated 15 times for each tube and the average was calculated. For each genotype 10 groups of females and 10 groups of males were tested. Flies were reared at 25°C and were maintained under 12hr light/dark cycle. Flies with an average age of 5 days were tested at RT, under the same light conditions. All assays were performed at the same time of day.

### Motor Behavior Analysis

Control and experimental flies were reared at 18°C and adult mated females up to 5 days post-eclosion were transferred to 25°C for behavioural analyses. Mechanical stimuli trains consisted of 5 pulses of 200ms each, spaced by 800ms. Motor behaviour parameters were determined as previously described (38,75,76).

### Bioinformatics Analyses and Identification of Cis-Regulatory Elements (CREs)

The Janelia Gal4 collection (http://flweb.janelia.org/cgi-bin/flew.cgi) (48) was screened for AMMC-specific GFP expression patterns. All hits were cross checked to be verified/excluded from the Janelia/Bloomington list (see https://bdsc.indiana.edu/stocks/gal4/gal4_janelia_info.html). For each hit, annotated left and right primers were used to BLAST the *Drosophila* genome annotated at the *Ensembl* genome browser (http://www.ensembl.org/index.html) to determine the position and sequence within genome version BDGP6, or where known the sequence was extracted from the respective gene map annotation in JBrowse (http://flybase.org/). The resulting sequence was compared against available VT enhancer sequences and their annotated expression pattern determined for DTB expression patterns (http://enhancers.starklab.org/) (50). *Drosophila*-specific, non-coding regulatory sequences were then used to BLAST the mouse (GRCm38.p5) and human (GRCh38.p7) genome annotated at Ensemble (https://www.ensembl.org/index.html) to identify any potential corresponding sequences. Any matching sequences were scrutinized for further analysis on the basis of criteria that have been used previously to identify transphyletic cis-regulatory DNA sequences (56). These criteria were:

i. the sequences are linked to the same homologous genes in the different species;
ii. there is a minimum of 62% sequence identity over at least 55 bp with minimum 1e^-1^ confidence level;
iii. the CREs are not un-annotated protein sequences; and
iv. the CREs are not repetitive elements.

The resulting sequences were then used for refined comparisons using pair-wise and multiple sequence alignment algorithms including EMBOSS Matcher and t-coffee (http://www.ebi.ac.uk/services). To carry out sequence alignments, which automatically indicated whether any CNS regulatory elements might be covered by the input sequences, we used the MLAGAN algorithm (http://genome.lbl.gov/vista/lagan/submit.shtml). Detected CNS CREs were then further scrutinized using the VISTA enhancer browser (https://enhancer.lbl.gov/frnt_page_n.shtml), which provides human and mouse regulatory sequences and their expression pattern at embryonic stage E11.5 in transgenic mouse embryos expressing LacZ under the control of the respective regulatory sequence (52,55). The relevant images of LacZ expression were extracted and reproduced with permission by Dr. Len Pennacchio, Lawrence Berkeley National Laboratory. Finally, identified MHB-specific regulatory sequences were utilized to perform multiple sequence alignment with the respective DTB->AMMC-specific regulatory elements; any matches were re-confirmed and quantified using the EMBOSS Matcher and CLUSTAL Omega (http://www.ebi.ac.uk/Tools/msa/clustalo/) algorithms.

### Statistical Analysis

Each data set was tested for normality using the Anderson-Darling test with α = 0.05. If every data set under comparison was normal and the variances were similar (Hartley’s fmax was calculated in each case and used as a cut-off for variance ratio), then a one-way ANOVA test was used to determine whether any differences existed between groups. If significance was found for ANOVA with α = 0.05, then pair-wise comparisons were made using a post hoc Tukey-Kramer test, again with α = 0.05. If any of the data sets was found not to be normally distributed, then a Kruskal-Wallis test was used to determine any overall differences between the groups with α = 0.05. If significance was achieved, a post hoc pairwise Mann-Whitney U test with Dunn-Sidak correction was used to compare groups with α = 0.05. For each test group, two controls were used corresponding to the two genetic elements that were altered in the group under analysis. For example, *R11A07>TNT* would be tested against TNT control and *R11A07>w1118*. For a result to be considered significant the experimental group had to be significantly different from both controls and the controls not to be significantly different from one another. All calculations were performed using MATLAB.

## Supporting information

Bridi_Supplementary Information

Movie S1. Startle-induced negative geotaxis of R52F05/w1118 control flies.

Movie S2. Startle-induced negative geotaxis assay of R52F05>TNT flies.

## SUPPLEMENTARY INFORMATION

Supplementary Information includes supplementary Figs. S1 to S9, Tables S1 to S2; captions for movies S1 to S2; captions for databases S1 to S3 and references for SI reference citations.

## ACKNOWLEDGEMENTS

We thank H. Karten, C. Ragsdale, A. Kamikouchi, A. Delogu, M. Fanto, and T.J. Weidemann for discussions; G.M. Rubin and L. Pennacchio for CRE data; D. Diaper and M. Dyson for graphics; and U. Walldorf, M. Noll, S. Carroll, A. Mueller, M. Michaelson, M. Landgraf, A. Ghysen, A. Gould, as well as the Developmental Studies Hybridoma Bank and the VDRC and Bloomington Stock Centres for providing antibodies and fly strains. This work was supported by a PhD fellowship from CAPES Foundation – Ministry of Education of Brazil to J.C.B. (BEX 13162/13-6); an IoPPN-King’s Independent Researcher Award to B.K.; an FWO post-doc grant to L.V.B.; the Flemish FWO (G065408.N10 and G078914N) to P.C.; the National Science Foundation under Grant No. 1754798 awarded to N.J.S.; and the UK Medical Research Council (G0701498; MR/L010666/1), the UK Biotechnology and Biological Sciences Research Council (BB/N001230/1) and the MND Association (Hirth/Nov15/914-793; Hirth/Oct13/6202; Hirth/Mar12/6085; Hirth/Oct07/6233) to F.H.

## AUTHOR CONTRIBUTIONS

J.C.B., B.H., Z.N.L., L.V.B. and B.K. performed experiments; F.H. and M.G. performed CRE analysis and phylogenetic comparisons; J.D. performed statistical tests for SING data. All authors analysed the data; F.H., P.C. and N.J.S. prepared the manuscript; F.H. conceived and designed the project.

## COMPETING FINANCIAL INTEREST

B.K. is co-founder of BFKLab LTD. The remaining authors declare no competing financial interest.

