## Supplementary material for "Ancestral Regulatory Mechanisms Specify Conserved Midbrain Circuitry in Arthropods and Vertebrates": Bridi_Supplementary Information

**This pdf file includes:**

Figs. S1 to S9
Tables S1 to S2
Captions for movies S1 to S2
Captions for databases S1 to S3
References for SI reference citations
Data Set S1 to S3

**Other Supplementary Information for this manuscript includes the following:**

Movies S1 and S2

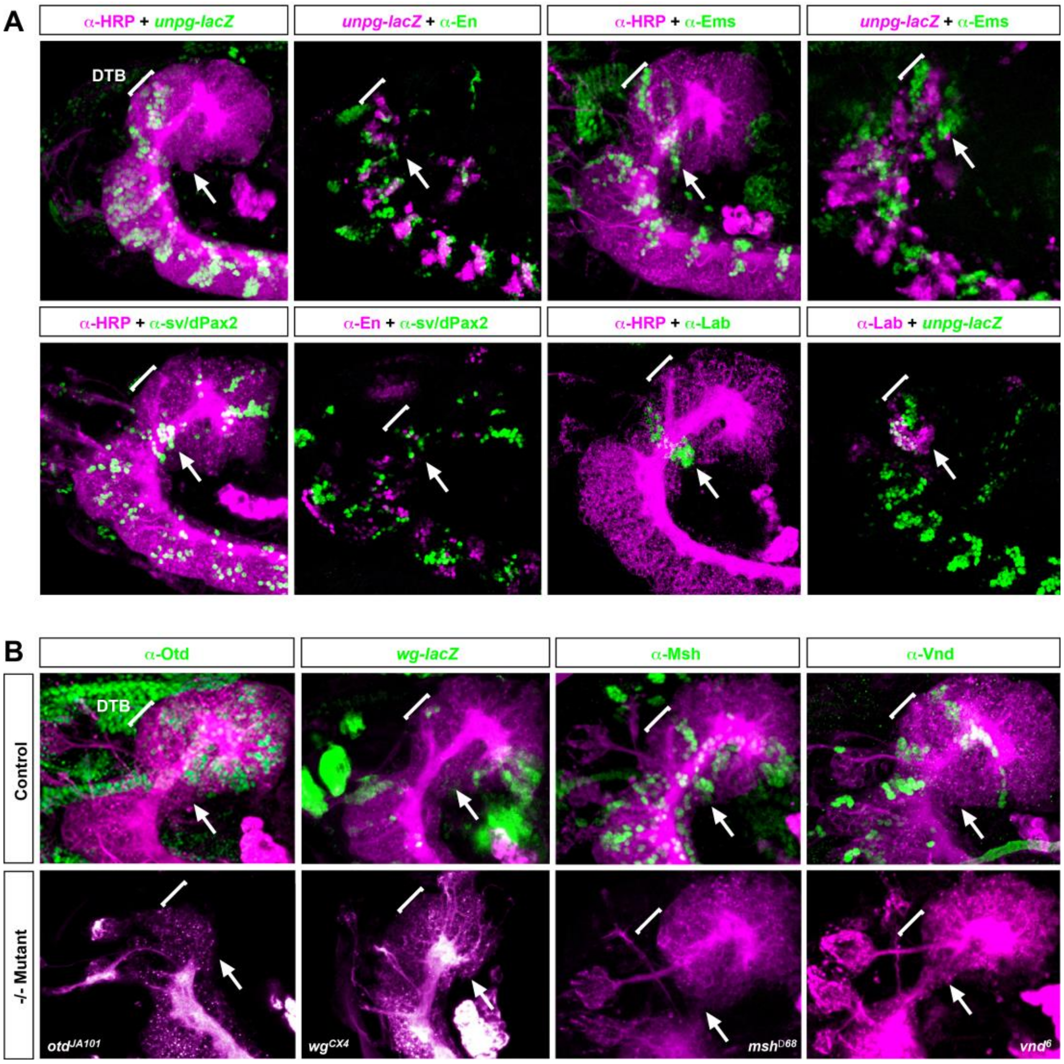


**Fig. S1.** Expression and function of genes that characterize the DTB region in the embryonic brain of *Drosophila.* (A) confocal images of stage 13-15 embryonic brains, lateral views; brackets indicate position of the deutocerebral-tritocerebral boundary (DTB) region, arrows highlight DTB-specific expression domains. From left to right, upper row: anterior-most expression of *Gbx* homologue *unplugged* (*unpg*) demarcates the DTB. *engrailed* (*en*) expression at the DTB coincides with *unpg* expression. Anterior-most expression of *empty spiracles* (*ems*) coincides with the DTB. Anterior-most expression domains of *unpg* and *ems* within the DTB. Lower row: prominent *sv/dPax2* expression domain within the DTB. DTB-specific expression of *sv/dPax2* coincides with *engrailed* expression at the DTB. The Hox1 ortholog *labial* (*lab*) demarcates the posterior part of the DTB and partially overlaps with the anterior *unpg* expression domain. (B) expression and function of genes involved in anterior-posterior (AP) and dorso-ventral (DV) axis specification. The *Otx* homologue *orthodenticle* (*otd*) posterior-most expression in the forebrain demarcates the DTB. HRP-labelled embryonic brain of *otd* null mutant reveals patterning defects, deleting all structures anterior to the tritocerebrum, including the DTB. Expression of *wingless* (*wg-lacZ*) in the anterior-most part of the embryonic brain and in a pattern coinciding with the DTB. HRP-labelled embryonic brain of *wg* null mutant reveals brain patterning severely affecting the DTB. Expression of *muscle-specific homeobox* (*msh*) in the anterior embryonic brain, including prominent DTB-specific expression pattern. HRP-labelled embryonic brain of *msh* null mutant reveals DTB patterning defects. Expression of *ventral nervous system defective* (*vnd*) in the anterior embryonic brain, including prominent DTB-specific expression pattern. HRP-labelled embryonic brain of *vnd* null mutant reveals severe DTB patterning defects.

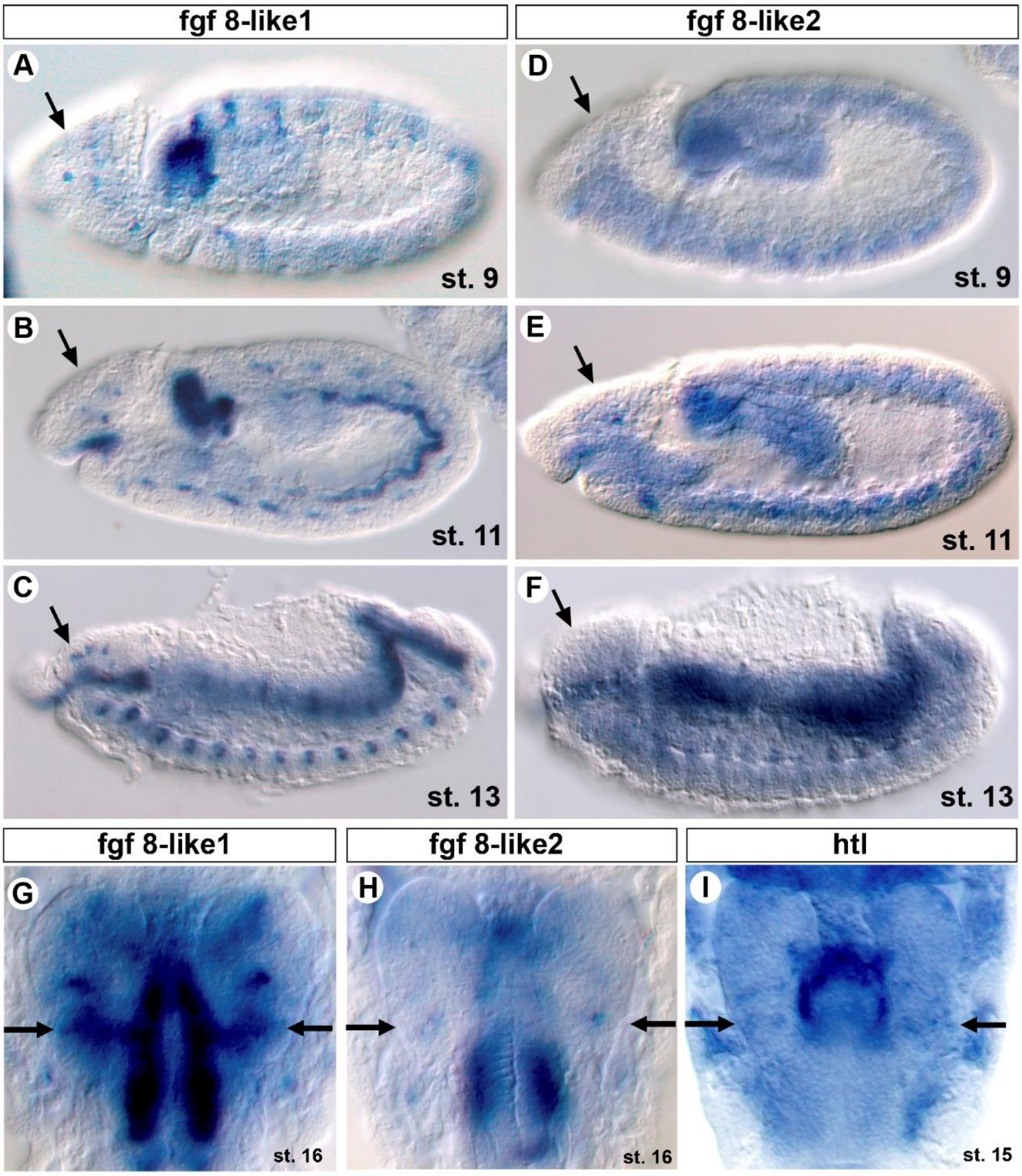


**Fig. S2.** Expression of *FGF8-like1* and *FGF8-like2* in the embryonic nervous system and brain of *Drosophila.* Whole-mount in situ hybridization of transcripts of (A-C) *FGF8-like1 (thisbe, ths)* and, (D-F) *FGF8-like2 (pyramus, pyr)* in wildtype embryos at indicated stages. Lateral views, anterior is to the left; arrows demarcate the primordium of the deutocerebral-tritocerebral boundary (DTB) region. (G, H) Anterior brains of stage 16 embryos, dorsal views; arrows demarcate *FGF8-like1* and *FGF8-like2* expression in the DTB. (I) In situ hybridization of FGF8-like receptor *heartless* (*htl*) transcripts in stage 16 anterior brain of wildtype embryo; arrow demarcates *htl* expression in the DTB.

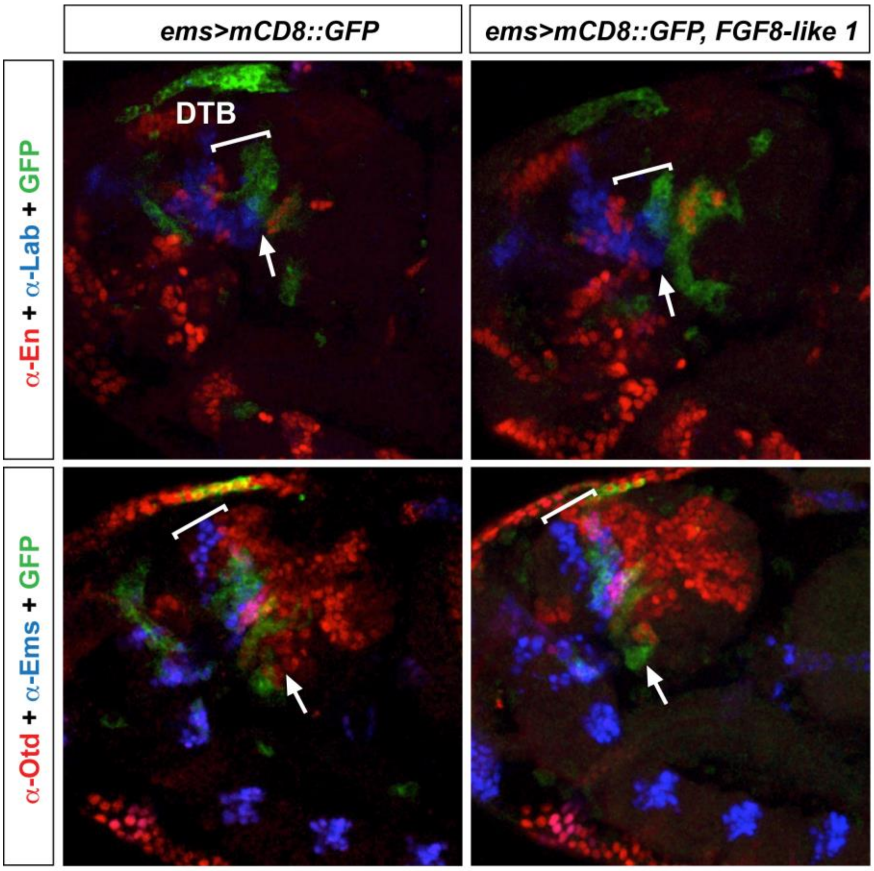


**Fig. S3.** FGF8 signalling in the embryonic DTB does not exert organiser activity. Ectopic expression of *FGF8-like1* does not affect DTB formation**.** Confocal images of developing embryonic brain of control (*ems-Gal4/UAS-mCD8::GFP*) and experimental flies (*ems-Gal4/UAS-mCD8::GFP; UAS-FGF8-like1)*; anterior is to the left. DTB-specific (bracket) expression patterns of *engrailed* and *labial*, as well as of *otd* and *ems* are unaltered in *ems-Gal4/UAS-mCD8::GFP; UAS-FGF8-like1* flies compared to controls (arrows).

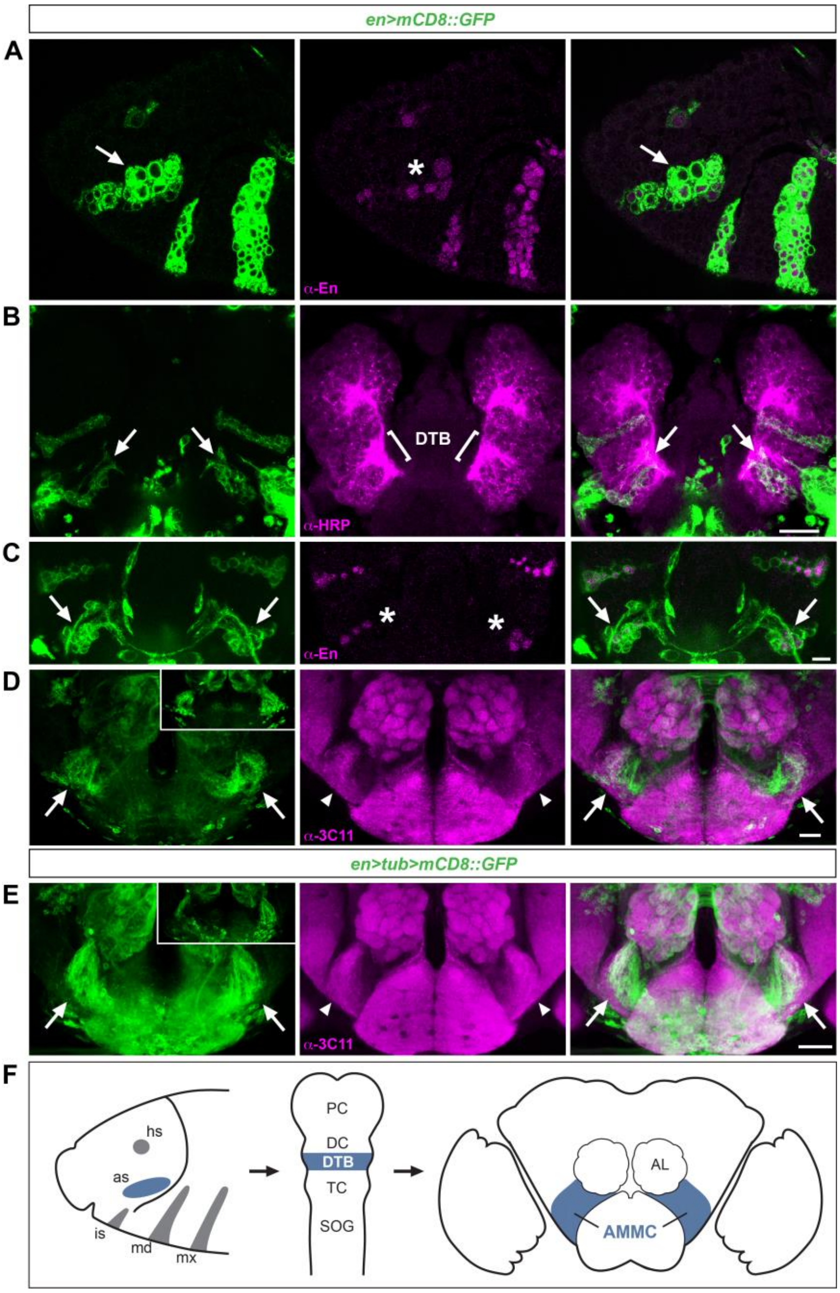


**Fig. S4.** The embryonic DTB region gives rise to neurons of the AMMC in the adult brain of *Drosophila.* Confocal images of developing embryonic (A-C) and adult brain (D, E); anterior to the left in A; and frontal in B-E. (A) at embryonic stage 11, *en-Gal4/UAS-mCD8::GFP* mediated cell labelling identifies anti-En positive neuroblasts in the procephalic neurectoderm, including NBs (asterisk) of the deutocerebral-tritocerebral neurectoderm boundary (DTB, arrows). (B) by embryonic stage 15, *en-Gal4/UAS-mCD8::GFP* identifies En-NB derived lineages, including DTB lineages (arrows) that (C) express Engrailed; their axonal projections include the developing antennal nerve (arrows). (D) in the adult brain, *en-Gal4/UAS-mCD8::GFP* visualises neurons, projections and axon terminals of the antennal mechanosensory motor centre (AMMC, arrows). (E) lineage tracing of *en* expressing central brain NB lineages. Confocal images of *en>tub>mCD8::GFP* identify AMMC neurons (inset) and axon terminals within the AMMC (arrows). (F) left, schematic of Engrailed expression domains in the procephalic neuroectoderm of stage 11 embryo, including head spot (hs), antennal spot (as) and intercalary spot (is); middle, schematic of stage 15 embryonic brain with protocerebrum (PC), deutocerebrum (DC), tritocerebrum (TC) and subesophageal ganglion (SOG); right, adult brain; AL, antennal lobe. Scale bars: 10μm (b), 50μm (D, E).

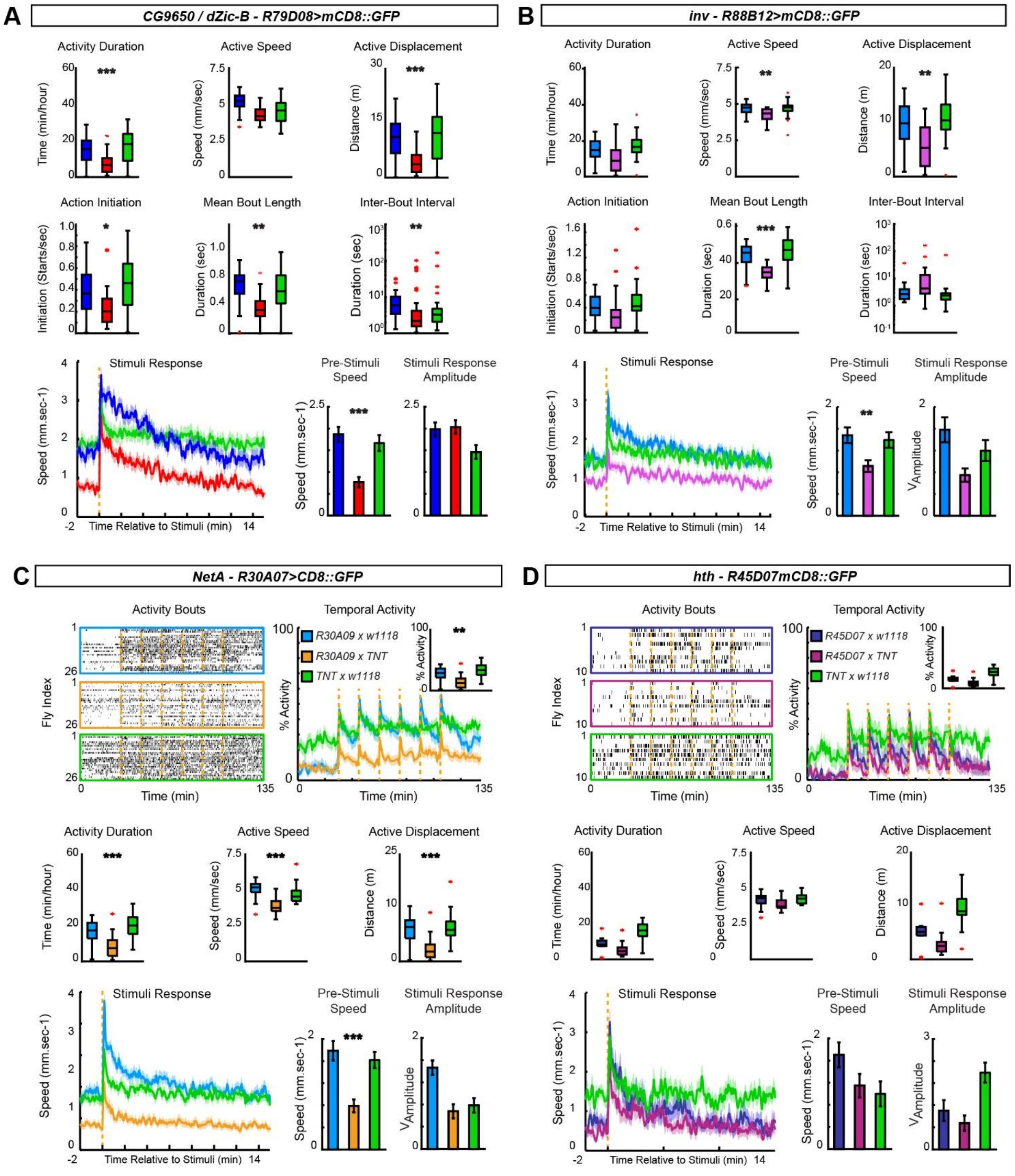


**Fig. S5.** Motor coordination in *Drosophila* is mediated by DTB-derived AMMC neurons. Motor kinematics and stimulus response (lower lane) of (A) *R79D08>TNT* and (B) *inv R88B12>TNT* with respective UAS/+ and Gal4/+ control flies; colour code indicate genotypes. Motor behaviour of (C) *R30A07>TNT* and (D) *R45D07>TNT* with respective UAS/+ and Gal4/+ control flies; top left, raster plots of activity bouts, each lane one individual fly; middle lane, motor kinematics; bottom lane, stimulus response. Mean ± Standard Error of the Mean (SEM), asterisks indicate $p<0.05$ (*), $p<0.01$ (**) or $p<0.001$ (***).

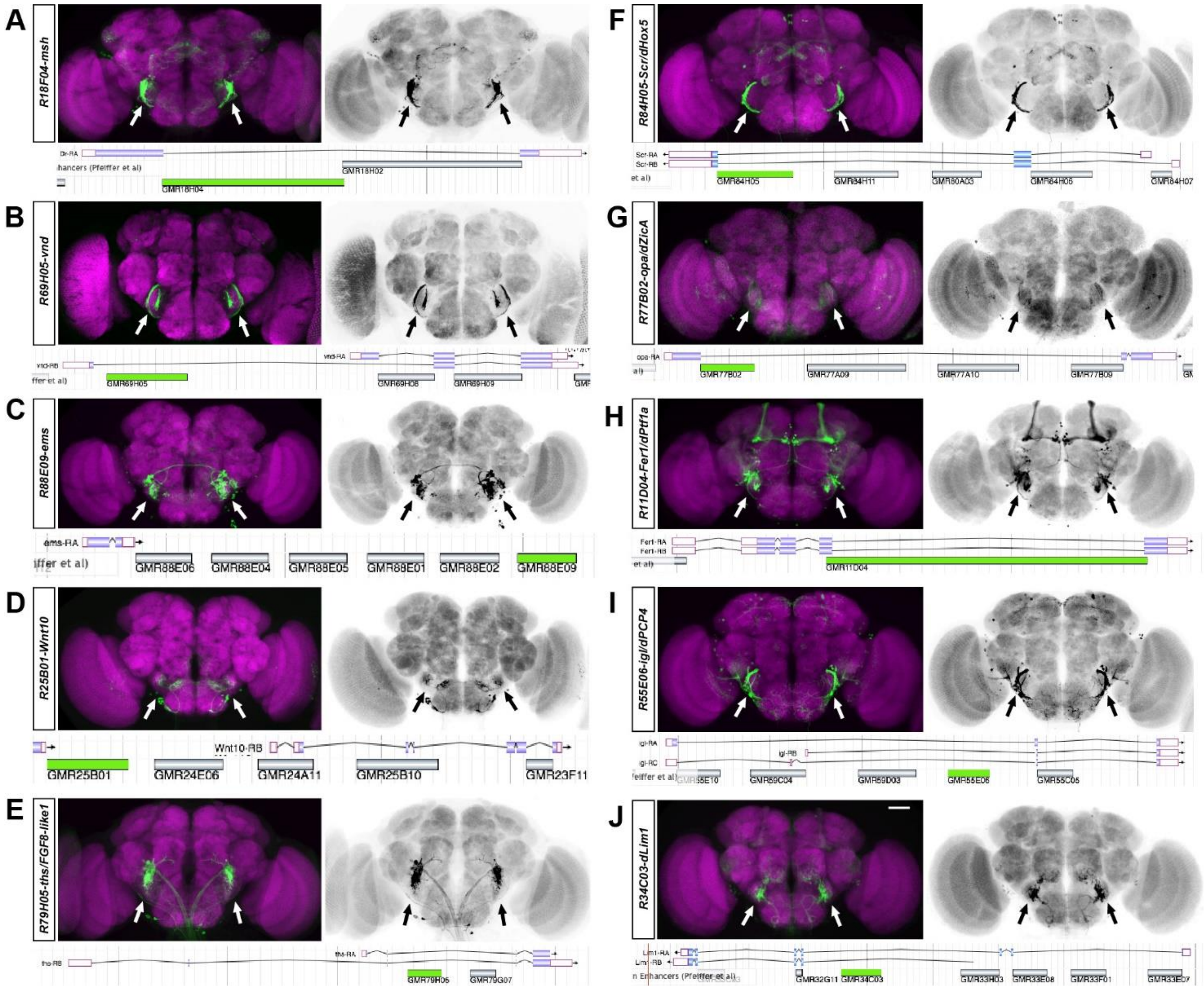


**Fig. S6.** Non-coding regulatory elements direct AMMC-specific expression in adult brain of *Drosophila.* Non-coding cis-regulatory elements (CREs) identified by the Janelia FlyLight project drive Gal4 expression of *UAS-mCD8::GFP* targeted to the antennal mechanosensory motor centre (AMMC). Confocal images of adult brains immunolabelled with anti-Brp/nc82 (magenta) and mCD8::GFP (green); black and white images are inversions to highlight GFP-labelled structures; dorsal is up (original confocal stacks are derived from the Janelia FlyLight repository). (A) *msh*-specific *R18F04>mCD8::GFP*. (B) *vnd*-specific *R69H05>mCD8::GFP*. (C) *ems*-specific *R88E09>mCD8::GFP.* (D) *Wnt10*-specific *R25B01>mCD8::GFP.* (E) *ths/FGF8-like1*-specific *R79H05>mCD8::GFP.* (F) *Scr*-specific *R84H05>mCD8::GFP.* (G) *opa/dZic4*-specific *R77B06>mCD8::GFP.* (H) *Fer1/dPtf1a*-specific *R11D04>mCD8::GFP.* (I) *igl/dPCP4*-specific *R55E06-Gal4/UAS-mCD8::GFP*. (J) *dLim1*-specific *R34C03>mCD8::GFP.* Arrows indicate GFP-labelled AMMC-specific neurons and/or projections/arborisations targeted by respective CRE. The genomic position of each CRE is depicted underneath the respective labelled brain and highlighted in green. Scale bar: 50μm.

*shaven - dPax2*

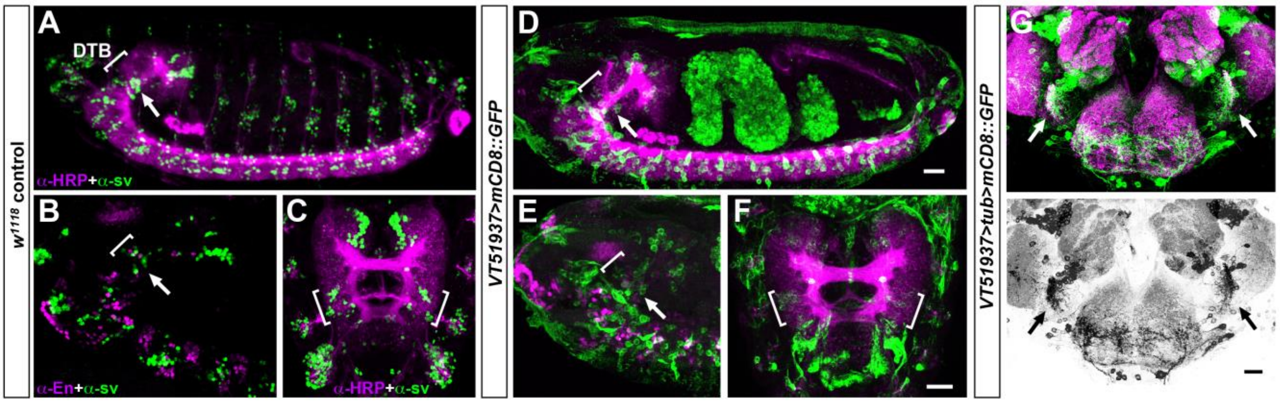


**H CRE sequence comparison**

```
shaven   CAACCAATTTTAGTT----TTCAG---CCCTCCACTCA------AATACATTGAAATGTCCA-TGAAGCGGGATGTACTTTT---CGCGCA----TCCCA
mPax2    CAAGCAATTTTAATATTTCTTCAGGGACTTTCCAGCAACAGCACAAGTCATTAATTCACCCTCCCAAATTGCTTATTCAATAGCAGGAACATGGCTCCTG
hPAX2    CAAGCAATTTTAATATTTCTTCAGGGACTTTCCAGCAACAGCACAAGTCATTAATTCACCCTCCCAAATTGCTTATTCAATAGCAGGAACATGGCTCCTG
         *** ******** *     *****    *  ****    *       **  **** *      **     **    *  * * * *    *  **     ***

shaven   AACATATGTCTCA--A-TTGTGTGCACTGTCGAAATATCCAATTGCTCATCCCG-TTTCAAGCT-GGTACTCAAAAAT--ATATAAAAC-CTGAAACTGT
mPax2    AATAA-AGTCCCAGAATTTATGTG-AC--TTGCACGAGTCAGGAGGTCAAACAACTGTTATGGAGCGAAGTTAAAAATCCAAATAAAATATTGACACTTT
hPAX2    AATAA-AGTCCCAGAATTTATGTG-AC--TCGCACGAGTCAGGAGGTCAAACAACTGTTATGGAGCGAAGTTAAAAATCCAAATAAAATATTGACACTTT
         ** *    *** **   * ** **** **   * * *   *  **    * ***  *    * * * *     * * * ******  * ******    *** *** *

shaven   TGTCTTAAGG-GCACAAACTAGGCAGAGTTAACTTCTGCCCTTACAACCCCCTGTAGCATATGGTAACCTTAACAATCTGTAAAAAAAACCTAAAAGATA
mPax2    TTTGGGGAGGGGAAGAGAGGAGGCCAAGGTATTT------AAAATAGGTTTTTGTTGCA-ATG----CCTC------CAGGATAAAGAAT-GGAAGGGGA
hPAX2    TTTGGGGAGGGGGAGGTGGGAAGCAAAGGTATTT------AAAATAGGTTTTTGTTGCA-ATG----CCCC------CAGGATAAAGAAT-GGAAGGAGA
         * *    *** * *       * **  ** **  *          * *       *** *** ***     **          * * * *** **     ** *  *

shaven   CTTAAATCAATGCTATCCCGGAGGTCCTATGAGTTTTAATAGGTGTATTTGGGTACCGCATTTTGTAATTATTAAAGTTTATTGCA-AA
mPax2    CTGAAGACAAG------TTGGAGTT--TATAAAA-TCAATGGAATTTATTGCACATC-TACTTGGCATTTAATAAT-TCTCTCATT-TC
hPAX2    CTGAAGACAAG------TTGGAGTT--TATAAAA-TCAATGGAATTTATTGCACAGC-TACTTGGCATTTAATAAT-TCTCTCATTTCC
         ** **  ***           **** *  *** *   * *** *   *  ***   * *  * ** * * *** ***  * * *
```

**I Intragenic location of CRE**

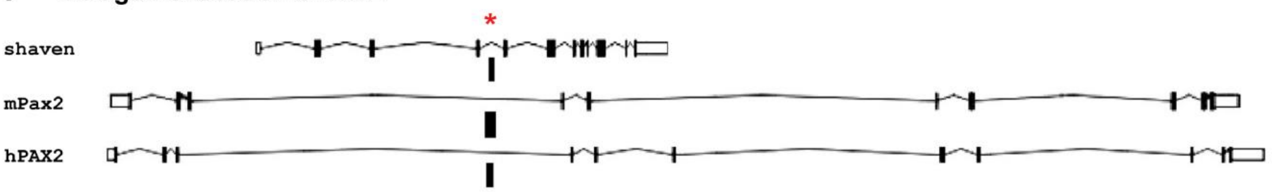


**Fig. S7.** Conserved non-coding regulatory sequence of *shaven/dPax2* directs DTB-AMMC specific expression in *Drosophila.* (A-C) confocal images of anti-sv/dPax2 immunolabelling (green, yellow) in embryonic CNS (A, lateral, anterior to left) and brain (B, lateral; C, frontal) co-immunolabelled with anti-HRP (A, C, magenta) or anti-Engrailed (B, magenta); brackets indicate deutocerebral-tritocerebral boundary (DTB) region; note sv/dPax2 expression within DTB (arrows). (D-F) lacZ expression directed by *sv/dPax2*-specific regulatory element VT51937, including DTB (brackets, arrows). (G) *sv/dPax2-VT51937>tub>mCD8::GFP* mediated genetic tracing identifies AMMC neurons and projections (arrows). (H) Comparison of non-coding regulatory sequences (CREs) of *Drosophila melanogaster sv/dPax2* that comprise parts of *VT51937*, mouse *mPax2* and human *hPAX2* (asterisks denote identical amino acids), and (I) their comparable intragenic locations (black bar, see arrow above exon-intron annotation) within genomic loci of, respectively, *sv/dPax2*, *mPax2* and *hPAX2.* Scale bars: 20μm.

##### *invected - ENGRAILED 2*

**A** **CRE Sequence comparison**

```
invected   TGAAAACTGAAGACTGCAACCAGGGACCACACACGACGATCGAGGCTGTGCCACAAGCTGCTCTGGGCAACGGAAGGCGGCAACATCGACAGTAGTTTTG
mEn2       TCAAACTTGCATA-TGCAACGAGGT-----GCTG--------------------GA---GCTTTGT---------GGCGGAAATAT-------AATTAAA
hEN2       TCAAACTTGAATA-TGCAATGAGAC-----GCGG--------------------GA---GCTTTGT---------GGCGGAAATAT-------AATTAAA
           * ***  ** * * *****  **        *                       *   *** **          ***** ** **       * **

invected   CGGCTCGTTTGCTGCGTTGACTCTTCGCCGAGTCCCGTGATCATGGACATGATCATCATGACGTACGATGATCGTGGGGTCCGCAGAACGTCATCATCGC
mEn2       -----------CTG-------------GTGAGA--------GATGTAAAAGA------TGA-------AGAA--TGGG---------ATGACATCAGA--
hEN2       -----------CTG-------------GGGAAA--------GATGTAAAAGC------TGA-------AGAA--TGGG---------ATGACATCAGG--
                      ***               **           *** * * *       ***        **   ****         * * *****

invected   GAAGATCTTCCTCCTCCTGATGCGGCGCCAAATTGCAGCAAAGTAAAACAGCGTTTAGCATTGTGCTTGTAATGATCTTGGTCAGGCA-ATCCGAGGGCC
mEn2       -CTGATTTCAC-ACT--TG--GCA--GTCCGATAGCTGGG--CCCAAGCCGGG------------CTTGCCACG-------CAAGGCAGATCAAAGTGCT
hEN2       -CTGATTTCAC-ACT--CG--GCA--GTCGGATCGCTGGG--CCCAAGCCGCGCT----------CTTGCCACG-------CAAGGCAGATCAAAGTGCC
              *** *  *  **   *  **   * *  ** ** *       ** * * *            ****  * *         ***** ***  ** **

invected   TGGCGTCCACAAATCTACAGAAGAACCCAAGTCTGGAGATTCAAAGAGAGGTCCCCGAGATTCTGGGCGACTTAAGAGAAGAACAGCGCAACTCGTTCGC
mEn2       CTGCCACCACTAA--------A-AAGGCAAA--GGGAGACTTAAGTA------TGCTA-ATC--------CCCAGGACAAATA-----------------
hEN2       CTGCCACCGCTAA--------A-AAGGCAAA--AGGGGACTTAAGTA------TGCTA-ATC--------CCCAGGACAAATA-----------------
             **  ** * **        * **  ***    ** ** * **  *        * * **         *  * ** **  *

invected   TGAGTTAGCGCCTTTGCAATGGGTGTAATAATTTATGTATTTTGATTAGGCCGAAGTTTCTGTGACTGTGGCTCGGCTTTTGCCTTTTAATTGATAAGTG
mEn2       ----------------------------TATTTTA---ATCTTGTTAGAATACAAGTTAATG---CTGCAGTGCAG---T-GGCT-GAACTTGGTCAGGC
hEN2       ----------------------------TATTTTA---ATCTTGTTAGAATACAAGTTAATG---CTGCAGCTCAG---T-GGCT-GAACTTGGTCAGAC
                                       ** ****   ** *** *       *****  **   ***  *  * *   * * **   * *** * **

invected   TGTTTGAGATACTTT-TTAATGAAGCTAATGAAACATGGTGAGCTTTCAGATATGTCTATATCTTTCAAGGGT-TTAGGCACTCCGGTAATTTAATAAAC
mEn2       AGG-AGAGACCCATTTTTATCTTAGCTA-TGAA------TGAGGT---GAA--------GCTCTT-----GGTTTTATTCA--TCAGTAATAAAA-GTAC
hEN2       TGT-AGAGATCCATTTTTATTTTAGCTA-TGAA------TGAGCT---AGA--------CCTCTT-----GGT-TTATTCA--TCGGTAATAAAA-GTAA
            *   ****  * ** ***    ***** ****      **** *     *          ****     *** ***  **   * *****  **   *

invected   ATTTAAATCTTTAAATGAA
mEn2       ATTTA-----TAAAACGAA
hEN2       ATTTA-----CAAAACGAA
           *****       *** ***
```

**B** **Intergenic location of CRE**

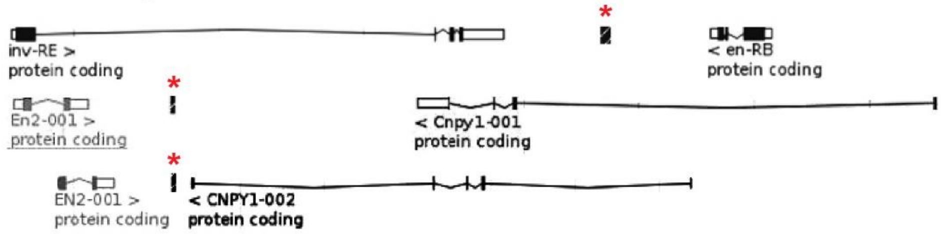


**C** **Hs CRE**

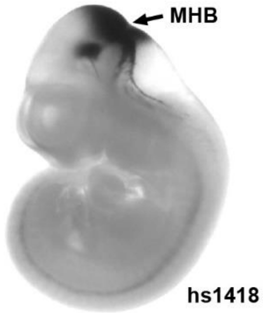


**Fig. S8.** Conserved non-coding regulatory sequence of *engrailed/invected* and *ENGRAILED 2* directs MHB-specific expression in mouse. (A) sequence comparison of non-coding cis regulatory elements (CREs) of *Drosophila melanogaster inv/dEn2*, mouse *Engrailed 2 (mEn2)* and human *ENRGAILED 2 (hEN2)* aligned towards consensus sequence with human CRE hs1418 reported in VISTA enhancer database (asterisks denote identical amino acids). (B) comparable intragenic CRE location (black bar) of, respectively, *Drosophila inv/dEn2*, mouse *mEn2* and human *hEN2*. Note inverted orientation of *mEn2* and *hEN2* compared to *Drosophila invected* and *engrailed.* (C) whole-mount E11.5 mouse embryo (VISTA database, reproduced with permission) showing LacZ expression pattern in the midbrain hindbrain boundary region (MHB, arrow) targeted by human EN2-specific CRE, *hs1418*.

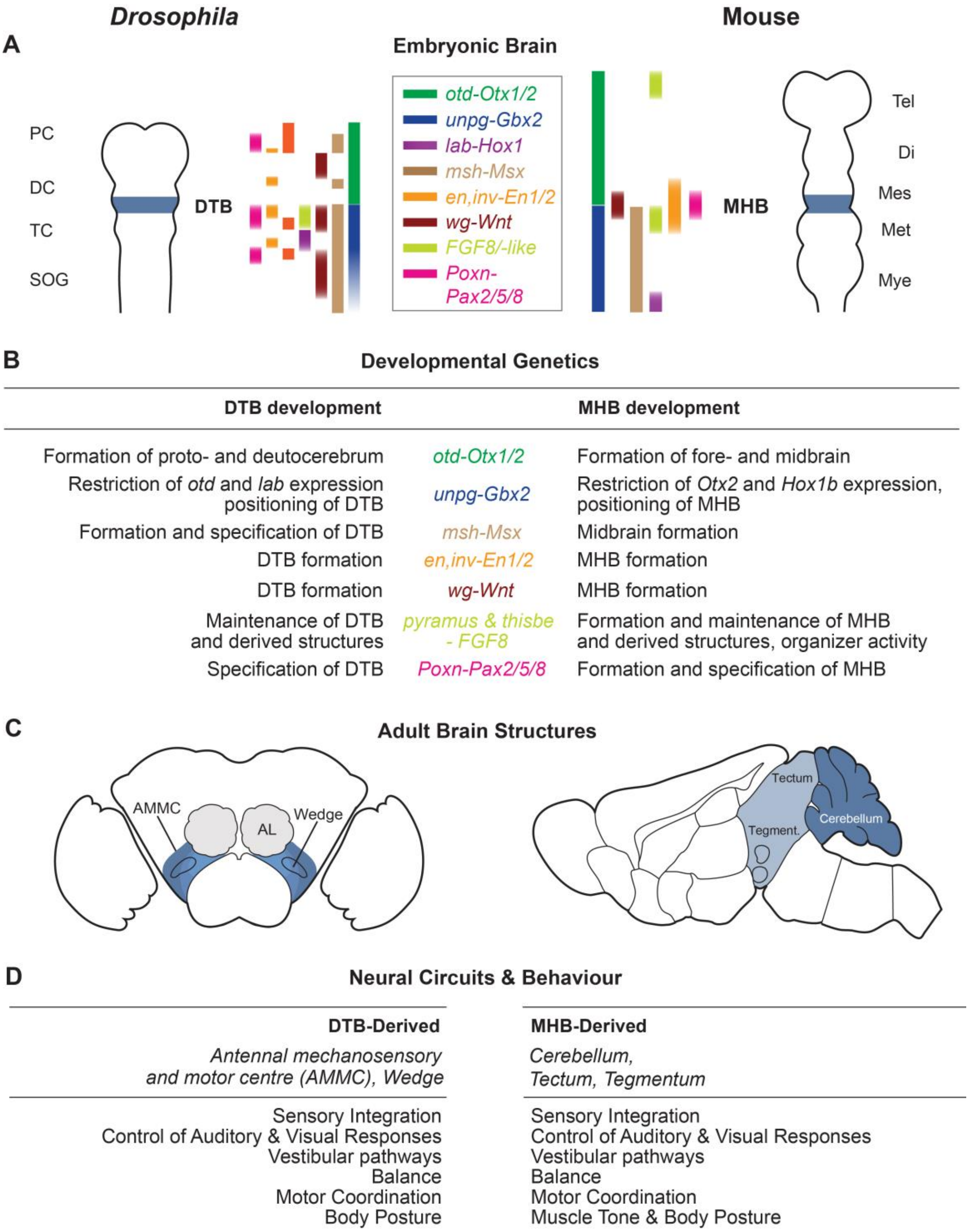


**Fig. S9.** Comparative analysis of ground pattern organization of the *Drosophila* DTB and the mouse MHB. (A) schematic of embryonic *Drosophila* and mouse brain showing expression patterns of homologous genes along neuraxes; anterior is to the top. (B) summary of DTB/MHB-related functions of gene homologues. (C) schematic of adult *Drosophila* and mouse brain; highlighted in blue are structures derived from, respectively, DTB and MHB regions. (D) DTB and MHB derived neural circuits in the adult brain and behavioural manifestations mediated by DTB/MHB-derived neural circuits. Abbreviations: AL, antennal lobe; AMMC, antennal mechanosensory motor center; dc, deutocerebrum; di, diencephalon; DTB, deutocerebral-tritocerebral boundary; hb, hindbrain; mb, midbrain; Mes, Mesencephalon; Met, Metencephalon; Mye, Myelencephalon; MHB, midbrain hindbrain boundary; pc, protocerebrum; SOG, Subesophageal ganglion; tc, tritocerebrum; tel, telencephalon.

**Table S1.** Statistical analysis of SING data

| | Genotype tested | Tests done (alpha=0.05)<br>KW = Kruskal-Wallis with post hoc Mann-Whitney U test<br><br>ANOVA = ANOVA with post hoc Tukey-Kramer | p value for group test (alpha = 0.05) | p values for pairwise tests<br><br>(alpha=0.05, Dunn-Sidak corrected value = 0.01695)<br><br>e.g. ctrl1 = TNT/w or nbc/w, ctrl2 = Gal4/w,<br><br>subject = Gal4>TNT<br><br>N/A signifies that no pairwise test was carried out because the group test did not show significant differences between groups | | | Significance<br><br>(requires p≤0.05 for group test AND ctrls not significantly different to one another AND subject significantly different to both controls) | Difference compared with controls |
|---|---|---|---|---|---|---|---|---|
| | | | | ctrl1 vs ctrl2 | subj vs ctrl1 | subj vs ctrl2 | | |
| All genotypes | *R11A07>TNT* | ANOVA | 0.0000 | Not Significant | Significant | Significant | Significant | Decrease |
| | *R37F03>TNT* | KW | 0.0000 | 0.0199 | 0.0021 | 0.0000 | Significant | Decrease |
| | *R52F05>TNT* | ANOVA | 0.0001 | Not Significant | Significant | Significant | Significant | Decrease |
| | *R51D11>TNT* | ANOVA | 0.0000 | Not Significant | Significant | Significant | Significant | Decrease |
| | *R79D08>TNT* | KW | 0.0000 | 0.6454 | 0.0000 | 0.0000 | Significant | Decrease |
| | *R70G01>TNT* | ANOVA | 0.0000 | Not Significant | Significant | Significant | Significant | Decrease |
| | *R24C06>TNT* | ANOVA | 0.0000 | Not Significant | Significant | Significant | Significant | Decrease |
| | *R45D07>TNT* | ANOVA | 0.0000 | Not Significant | Significant | Significant | Significant | Decrease |
| | *R22B11>TNT* | KW | 0.0000 | 0.1594 | 0.0000 | 0.0000 | Significant | Decrease |
| | *R55C02>TNT* | ANOVA | 0.0006 | Not Significant | Significant | Significant | Significant | Decrease |
| | *R87G01>TNT* | KW | 0.0016 | 0.4323 | 0.0047 | 0.0012 | Significant | Decrease |
| | *R30A07>TNT* | ANOVA | 0.0000 | Not Significant | Significant | Significant | Significant | Decrease |
| | *R19E09>TNT* | KW | 0.0000 | 0.0166 | 0.0000 | 0.0000 | Not Significant | No difference |
| | *R25B01>TNT* | KW | 0.0000 | 0.0222 | 0.0000 | 0.0000 | Significant | Decrease |
| | *R17A10>TNT* | KW | 0.0007 | 0.0498 | 0.0123 | 0.0006 | Significant | Decrease |

**Table S2.** *Drosophila* genes, their vertebrate homologs and role in the development of the cerebellum and cerebellar neural circuits.

| *Drosophila* | Vertebrate | Ref. | Embryonic | Postnatal |
|---|---|---|---|---|
| *orthodenticle* | *Otx1* | (1) | - | EGL |
| | *Otx2* | (1) | Cerebellar precursor cells | EGL and IGL |
| *unplugged* | *Gbx2* | (2) | MHB | - |
| *vnd* | *Nkx6.1* | (3) | MHB | ? |
| *msh* | *Msx2* | (4) | MHB | IGL, Purkinje cells |
| *engrailed/invected* | *En-1/En-2* | (5) | MHB | Granule cells |
| *wingless* | *Wnt-1* | (5) | Rostral to MHB | |
| | *Wnt-3* | (5) | MHB | Purkinje cells |
| | *Wnt-7a* | (5) | uniform brain expr. | Granule cells |
| *pyramus/thisbe* | *FGF8* | (6) | MHB | Cerebellum |
| *shaven/dPax2* | *Pax2* | (5) | MHB | Cerebellum |
| *Poxn* | *Pax2/5/8* | (5) | MHB | Granule cells |
| *Enhancer of split* | *HES1/HES-3* | (7) | MHB | Purkinje cells |
| *Eyeless* | *Pax6* | (8) | Cerebellum | EGL, granule cells |
| *Odd-paired* | *Zic1/2* | (9) | Granule cell precursor | Granule cells |
| *CG9650* | *Zic1/2* | (9) | Granule cell precursor | Granule cells |
| *atonal/cato* | *Math1* | (10) | EGL | Granule cells |
| | *NeuroD* | (11) | EGL | IGL, granule cells |
| *Sex combs reduced* | *HoxA5* | (12) | Hindbrain | Purkinje cells |
| *Fer1* | *Ptf1a* | (13) | Cerebellum | GABAergic neurons |
| *igloo* | *PCP4* | (14) | Cerebellum | Purkinje cells |

Modified and updated after (15-17). Abbreviations: EGL, external granule cell layer; IGL, internal granule cell layer; MHB, midbrain hindbrain boundary; Ref., reference.

**Movie S1.** Startle-induced negative geotaxis of *R52F05/w*$^{1118}$ control flies.

**Movie S2.** Startle-induced negative geotaxis assay of *R52F05>TNT* flies.

**Additional data set 1 (separate file).** Super-conserved CRE sequences for *shaven/PAX2*.

**Additional data set 2 (separate file).** Super-conserved CRE sequences for *invected/engrailed* and *EN2*.

**Additional data set 3 (separate file).** Super-conserved CRE sequences for *dachshund/DACH1*.

### SUPPLEMENTARY DATA SET S1

#### *shaven/PAX2* superconserved sequence

***sv/dPax2* VT51937**

```
caacatcatcgtagaattttggccgtgccacgtggcagttcctaagtcgacactgtcccacattgctattatttacgcgcatccacttt
gaccataatttatgttggtagcagtcaacgaggccaacgagagcagctggaaaacctttaatcccatgttttatttgaaattagctgaa
ccctgtttaaatgtaaattttatgaaattttaaaaatgttaaagcccaaatagccgaacgtcctaaaaacatgtaattacgaaggtttt
tcttcaatcttacgattagtcgttttggaactatggaatatggttgtcccgtgttactcgtacctgttatggctttcaaacaagactag
tttggttaggaacgaatatggacccttattataaaatttttcgaagaataatgttacaaaatgttctttatatttaacaaaaagcata
caagtcattatctgttgatcgagtatttcaaatgtcaagtctcatctctattctgcatgctagaaaatgtaattttaattttaatttt
aaccgatctcaacacgtgcagttataatcaattcgattcgtgttaccgccgggtcccgcaaaaagtaacaaacgtctagattggcatgt
ggccgctggtaatgcaaaagttgcgacggacaagggtaattcatggtataatcaaggaggaagaaaaatttcataatgcgacaatgtgt
gatgtgatggcacttgaacgatgattagggtggcggtgattcaatcgaataaaaattgtgcgtacggcggtgccaggactccccttata
ctgattcccaggagactgtcacttaataaagtgactttcatagcactcaaccaatttagttttcagccctccactcaaatacattgaa
atgtccatgaagcgggatgtacttttcgcgcatcccaaacatatgtctcaattgtgtgcactgtcgaaatatccaattgctcatcccgt
ttcaagctggtactcaaaaatatataaaacctgaaactgttgtcttaagggcacaaactaggcagagttaacttctgcccttacaaccc
cctgtagcatatggtaaccttaacaatctgtaaaaaaaacctaaaagatacttaaatcaatgctatcccggaggtcctatgagttttaa
taggtgtatttgggtaccgcattttgtaattattaaagtttattgcaaaattgtacacaaatactaacacgacacgtttggcatctctt
gaatatgcagggtgtgtgctggacagaaggacaagccatgtccatgtatggactgaccaatattatttatatatctatattgcctcggg
aacgctttttacttgttacaacgaattttactttctataacattttattatcgtattagtataaaataataatacgataatattacga
ttacgattagtacgatctttaaaaacttattacatagttttgatttatatattttgaccacaaaaggcctgtaagacttagtagagcca
ccaaaactttaagtataagtactgaagtcataattagagattataaataacaatgatttacctacctaccctcatgggtaggtgatttt
aagtaatatgaacgttaaagtgttcatggctaacctggggaacagtcacttcttggatatctccatcttccttgcaagaagtgcaaggg
gaaccatagtggtctatcctcttttaaatattatatgtatgtacaactgattgtttattgtagttagttgtgtacaattgatttgttga
taagtatgtaaaaagtagaagaaaacgttctgacccaataaagttcttataatcttggttaggatcacttgccgagtgattgtggccat
gtccaaaactgttaaatatataattcttttctaactttattacaagactatttctgatccttcgctaacaaaatgttgaaattgggtaa
gggggatctgatagagactaggaactattgcggttcctattatttttagtaggcttaaataaaatttaccatttcccttctaaccgtg
aagagcgttgattttatttaactaagtcacttaaaatcttttcacttaatttcagaattgtaaggaataaagccgccgagaaagccaaa
cacgtacatcatcaccagcagcatcatgtttctcagagtctgggtgggggcatattgccacggaaagtgttgacagcagcacagga
```

*Green highlighted sequence depicts sv/Pax2 superconserved sequence*

**<u>EMBOSS MATCHER sv superconserved against VT51937</u>**

```
sv_CNS_CRE1        1 CAACCAATTTTAGTTTTCAGCCCTCCACTCAAATACATTGAAATGTCCAT     50
                     ||||||||||||||||||||||||||||||||||||||||||||||||||
sv_VT51937       849 CAACCAATTTTAGTTTTCAGCCCTCCACTCAAATACATTGAAATGTCCAT    898

sv_CNS_CRE1       51 GAAGCGGGATGTACTTTTCGCGCATCCCAAACATATGTCTCAATTGTGTG    100
                     ||||||||||||||||||||||||||||||||||||||||||||||||||
sv_VT51937       899 GAAGCGGGATGTACTTTTCGCGCATCCCAAACATATGTCTCAATTGTGTG    948

sv_CNS_CRE1      101 CACTGTCGAAATATCCAATTGCTCATCCCGTTTCAAGCTGGTACTCAAAA    150
                     ||||||||||||||||||||||||||||||||||||||||||||||||||
sv_VT51937       949 CACTGTCGAAATATCCAATTGCTCATCCCGTTTCAAGCTGGTACTCAAAA    998

sv_CNS_CRE1      151 ATATATAAAACCTGAAACTGTTGTCTTAAGGGCACAAACTAGGCAGAGTT    200
                     ||||||||||||||||||||||||||||||||||||||||||||||||||
sv_VT51937       999 ATATATAAAACCTGAAACTGTTGTCTTAAGGGCACAAACTAGGCAGAGTT   1048

sv_CNS_CRE1      201 AACTTCTGCCCTTACAACCCCCTGTAGCATATGGTAACCTTAACAATCTG    250
                     ||||||||||||||||||||||||||||||||||||||||||||||||||
sv_VT51937      1049 AACTTCTGCCCTTACAACCCCCTGTAGCATATGGTAACCTTAACAATCTG   1098

sv_CNS_CRE1      251 TAAAAAAAACCTAAAAGATACTTAAATCAATGCTATCCCGGAGGTCCTAT    300
                     ||||||||||||||||||||||||||||||||||||||||||||||||||
sv_VT51937      1099 TAAAAAAAACCTAAAAGATACTTAAATCAATGCTATCCCGGAGGTCCTAT   1148

sv_CNS_CRE1      301 GAGTTTTAATAGGTGTATTTGGGTACCGCATTTTGTAATTATTAAAGTTT    350
                     ||||||||||||||||||||||||||||||||||||||||||||||||||
sv_VT51937      1149 GAGTTTTAATAGGTGTATTTGGGTACCGCATTTTGTAATTATTAAAGTTT   1198

sv_CNS_CRE1      351 ATTGCAAA    358
                     ||||||||
sv_VT51937      1199 ATTGCAAA   1206

Identity: 358/358 (100.0%) - # Similarity:   358/358 (100.0%)
Length: 358 - Gaps: 0/358 (0.0%) - Score: 1790
```

**Human PAX2 - GRCh38 10:100767614-100768004**

CAAGCAATTTTAATATTTCTTCAGGGACTTTCCAGCAACAGCACAAGTCATTAATTCACCCTCCCAAATTGCTTATTCAATAGCAGGAA
CATGGCTCCTGAATAAAGTCCCAGAATTTATGTGACTCGCACGAGTCAGGAGGTCAAACAACTGTTATGGAGCGAAGTTAAAAATCCAA
ATAAAATATTGACACTTTTTTGGGGAGGGGGAGGTGGGAAGCAAAGGTATTTAAAATAGGTTTTTGTTGCAATGCCCCCAGGATAAAGA
ATGGAAGGAGACTGAAGACAAGTTGGAGTTTATAAAATCAATGGAATTTATTGCACAGCTACTTGGCATTTAATAATTCTCTCATTTCC

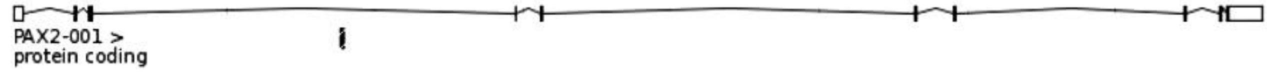


**Mouse Pax2 - GRCm38 19:44778358-44778747**

CAAGCAATTTTAATATTTCTTCAGGGACTTTCCAGCAACAGCACAAGTCATTAATTCACCCTCCCAAATTGCTTATTCAATAGCAGGAA
CATGGCTCCTGAATAAAGTCCCAGAATTTATGTGACTTGCACGAGTCAGGAGGTCAAACAACTGTTATGGAGCGAAGTTAAAAATCCAA
ATAAAATATTGACACTTTTTTGGGGAGGGGAAGAGAGGAGGCCAAGGTATTTAAAATAGGTTTTTGTTGCAATGCCTCCAGGATAAAGA
ATGGAAGGGGACTGAAGACAAGTTGGAGTTTATAAAATCAATGGAATTTATTGCACATCTACTTGGCATTTAATAATTCTCTCATTTC

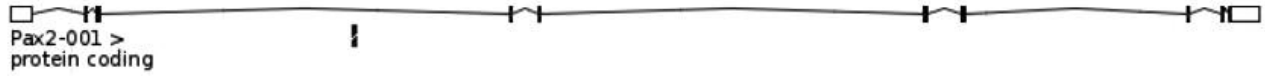


**Chicken scaffold >chromosome:Gallus gallus-5.0:6:17159089:17159892:-1**

CCTGGCACTGGAAGGGACAGCAAGGATGGAAGGCTGCTGTCAGCTGCACGGTGTGTGCGCAGGAATTCCCCTTCCTCTCCCAAGTGCGG
CTGTTTGTAGCCATTAAAAAGCCCAGGTCCTCATGCTGGGTGTTCAGTGAGTATTGATTCAGTCTG**CAAGCAATTTTAATATTTCTTCA**
**GGGACTTTCCAGCAACAGCACAAGTCATTAATTCACCCTCCCAAATTGCTTATTCAATAGCAGGAACATGGCTCCTGAATAAAGTCCCA**
**GAATTTATGTGACTCGCATGAGTCAGGAGGTCAAACAACTGTTATGGAGCGAAGTTAAAAATCCAAATAAAATATTGACACTTTTTTGG**
**GGA**

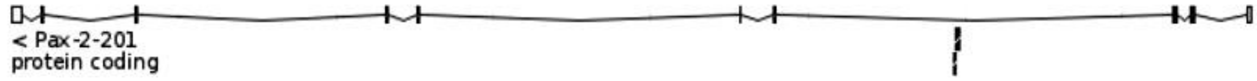


**Zebrafish Pax2a - GRCz10 13:29665906-29666048**

GCACATGCAAATATTAAAATCCACAAATCTAATCTAAAATGGCTTTTTAAGTTCATATAGATGGTTTGTGAAATTGTTTTATTATTATT
ATTATTATTATTAATTTATTGTATTTAATTTTACATTAGTA

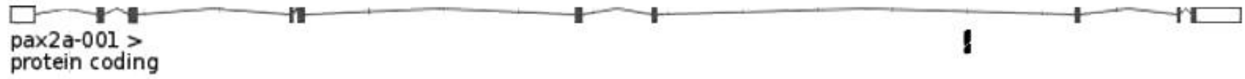


**MLAGAN comparison fly-mouse-human sv/Pax2**

| | |
|---|---|
| sequence1 | FlyShavenBDGP6_4_1083735_1116126_1:1-32392 (+) |
| sequence2 | mousePax2_GRCm38_19_44769940_44788670_1:1-18731 (+) |
| sequence3 | humanPAX2_GRCh38_10_100759566_100779388_1:1-19823 (+) |

Conservation ☑ Visible

ATGC CNS

```
sequence1(+)   000021153   TTCATAGCACTCAACCAATTTTAGTTTTCAGCCCTCCACTCAAATACATT   000021202
sequence2(+)   000008430   ----CAGTCTGCAAGCAATTTTAATATT----TCTTCA------------   000008459
sequence3(+)   000008060   ----CAGTCTGCAAGCAATTTTAATATT----TCTTCA------------   000008089

sequence1(+)   000021203   GAAATGTCCATGAAGCGGGATGTACTTTTCGCGCATCCCAAACATATGTC   000021252
sequence2(+)   000008460   ----------------GGGA-----CTTTCCAGCAAC---AGCACAAGTC   000008485
sequence3(+)   000008090   ----------------GGGA-----CTTTCCAGCAAC---AGCACAAGTC   000008115

sequence1(+)   000021253   -TCAATTGTGTGCACTGTCGAAATATCCAATTGCTCATCCCGTTTCAAG-   000021300
sequence2(+)   000008486   ATTAATT-----CACCCTC------CCAAATTGCTTATTCAATAGCAGGA   000008524
sequence3(+)   000008116   ATTAATT-----CACCCTC------CCAAATTGCTTATTCAATAGCAGGA   000008154

sequence1(+)   000021301   --CTGGTACTCAAAAATATATAAAACCTGAAACTGTTGTCTTAAGGGCAC   000021348
sequence2(+)   000008525   ACATGGCTCCTGA------ATAAAGTCCCAGAAT-TTATGTGACTTGCAC   000008567
sequence3(+)   000008155   ACATGGCTCCTGA------ATAAAGTCCCAGAAT-TTATGTGACTCGCAC   000008197

sequence1(+)   000021349   AAACTAGGCAGAGTTAACTTCTGCCCTTACAACCCCCTGTAGCATATGGT   000021398
sequence2(+)   000008568   GAGTCAGG--AGGTCAA-----------ACAAC------TGTTATGGAGC   000008598
sequence3(+)   000008198   GAGTCAGG--AGGTCAA-----------ACAAC------TGTTATGGAGC   000008228

sequence1(+)   000021399   AACCTTAACAATCTGTAAAAAAAACCTA----------------------   000021426
sequence2(+)   000008599   GAAGTTAAAAATCCAAATAAAATATTGACACTTTTTTGGGGAGGGGAAGA   000008648
sequence3(+)   000008229   GAAGTTAAAAATCCAAATAAAATATTGACACTTTTTTGGGGAGGGGGAGG   000008278

sequence1(+)   000021427   --------AAAGATACTTAAATCAA--TGCTATCCCGGAGGTCCTATGAG   000021466
sequence2(+)   000008649   GAGGAGGCCAAGGTATTTAAAATAGGTTTTTGTTGCAATGCCTCCAGGA-   000008697
sequence3(+)   000008279   TGGGAAGCAAAGGTATTTAAAATAGGTTTTTGTTGCAATGCCCCCAGGA-   000008327

sequence1(+)   000021467   TTTTAATAGGTGTATTTGGGTACCGCA-------------TTTTGTAATT   000021503
sequence2(+)   000008698   ---TAAAGAATGGA--AGGGGACTGAAGACAAGTTGGAGTTTATAAAATC   000008742
sequence3(+)   000008328   ---TAAAGAATGGA--AGGAGACTGAAGACAAGTTGGAGTTTATAAAATC   000008372

sequence1(+)   000021504   ATTAAAGTTTATTGCAAAATT-------GTACACAAATACTAACA-CGAC   000021545
sequence2(+)   000008743   AATGGAATTTATTGCACATCTACTTGGCATTTAATAATTCTCTCATTTCC   000008792
sequence3(+)   000008373   AATGGAATTTATTGCACAGCTACTTGGCATTTAATAATTCTCTCATTTCC   000008422
```

***D. melanogaster shaven* gene locus and homology to other insects**

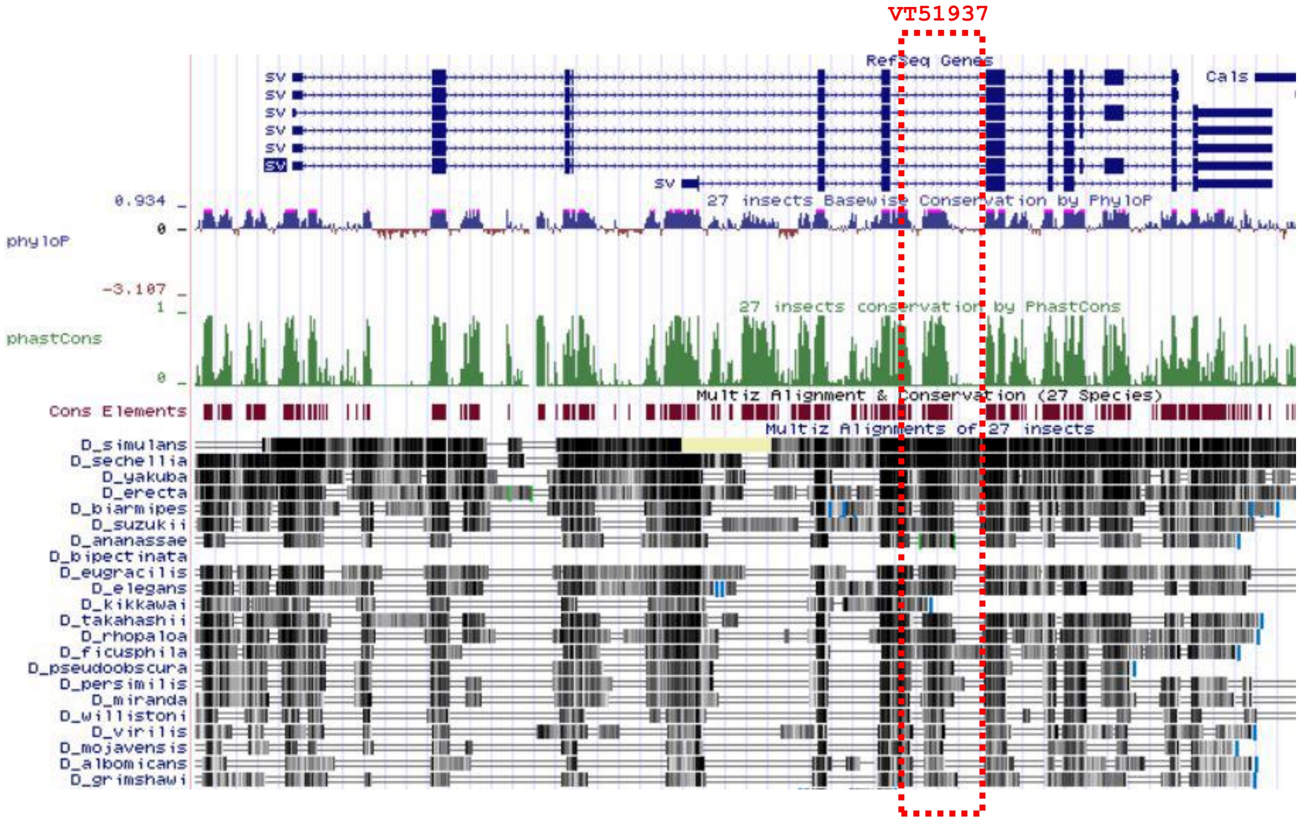


Image capture of UCSC Genome Browser on *D. melanogaster* Aug. 2014 (BDGP Release 6 + ISO1 MT/dm6) assembly, shown is the *shaven* gene locus. RefSeq Genes shows exon (blue bars) and intron structure; in red is shown Ensembl Gene annotation used for genome browsing. Green bar diagram indicates conservation among *Drosophilidae* species which are listed further below (black bars indicate sequence homology). Conserved elements are shown in dark red. Dashed red box indicates topology and extension of VT51937 CRE sequence determined in *Drosophila melanogaster*.

**Examples of sv/Pax2 superconserved sequence found in other *Drosophilidae***

**D simulans scaffold**
>chromosome:GCA_000259055.1:4:863940:864789:1
**CAACCAATTTTAGTTTTCAGCCCTCCACTCAAATACATTGAAATGTCCATGAAGCGGGATGTACTTTTCTCGCATCCCAA
ACATATGTCTCAATTGTGTGCACTGTCCAAATATCCAATTGCTCATCCCGTTTCAAGCTGGTACTCAAAAAAACATAAAA
CCTGAAACTGTTGTCTTCAGGGCACAAACTAGGCAGAGTTAACTTCTGCCCTTACAACCCCCTGTAGCATATGGTTACCT
TAACAATCTG**

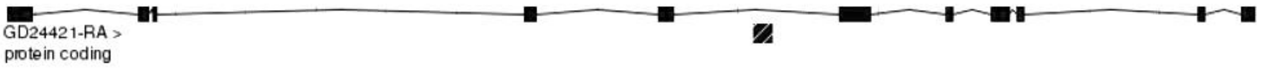


**D sechellia scaffold**
>supercontig:GCA_000005215.1:scaffold_40:169522:170371:1
**CAACCAATTTTAGTTTTCAGCCCTCCACTCAAATACATTGAAATGTCCATGAAGCGGGATGTACTTTTCTCGCATCCCAA
ACATATGTCTCAATTGTGTGCACTGTCCAAATATTCAATTGCTCATCCCGTTTCAAGCTGGTACTCAAAAAAACATAAAA
CCTGAAACTGTTGTCTTCAGGGCACAAACTAGGCAGAGTTAACTTCTGCCCTTACAACCCCCTGTAGCATATGGTTACCT
TAACAATCTG**

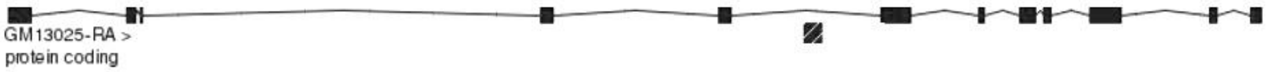


**D yakuba scaffold**
>chromosome:GCA_000005975.1:4:1205049:1205838:1
**CAACCAATTTTAGTTTTCAGCCCTCCACTCAAATACATTGAAATGTGCATG**T**GCGGGATGTACTTTTCTCGCATCCCAAA
CATATGTCTCAATTGTGTGCACTGTCCATATATCCAATTGCTCATCCCGTTTCAAGCTAGTACTCAAAAAAAAAGAAAAC**

**CTAAAACTGTTGTCTTCAGGACACAAACTGGGCAGAATTAACTTCTGCCCTTACAACCCTCTGTAGCATATGGTTACCTT AA**

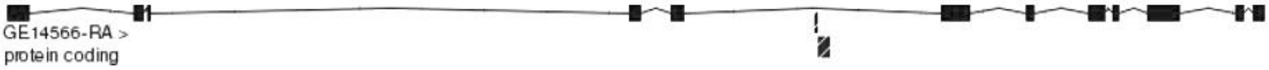


**D_pseudoobscura_scaffold**

>supercontig:GCA_000001765.2:Unknown_group_58:6806:7477:-1
**TGTACTTTACTCACATCCCAAACATATGTCTCAATTGTGTGCACTGCCCAAATACCCAATTGCTCATCCCGT**CTCTGAGC CGGCGCTTGCAGGAAAGAATCGAAAATCTGAAACTGATGTCAGCTTCCATCAT**TTGGACTGAATGGGCTTCTCTTCTT**

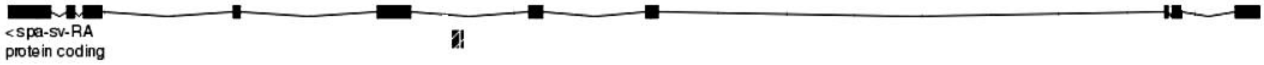


**D_ananassae**

**AACATATGTCTCAATTGTGTGCACTGCCCAAATATCAAATTGC**

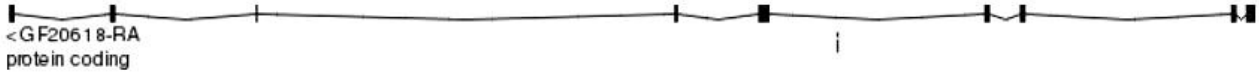


**D_willistoni**

>supercontig:GCA_000005925.1:scf2_1100000004943:1805136:1805769:-1
**CGCATCCCAAACATATGTTTCAATTGTGTTCACT**

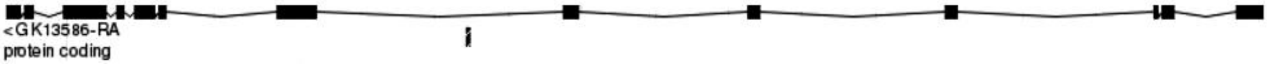

#### SUPPLEMENTARY DATA SET S2

#### *invected/engrailed - EN2* superconserved sequence

**Fly inv/en superconserved**

```
Location: BDGP6:2R:11516158:11517373:1
TGAAAACTGAAGACTGCAACCAGGGACCACACACGACGATCGAGGCTGTGCCACAAGCTGCTCTGGGCAACGGAAGGCGGCAACATCGA
CAGTAGTTTTGCGGCTCGTTTGCTGCGTTGACTCTTCGCCGAGTCCCGTGATCATGGACATGATCATCATGACGTACGATGATCGTGGG
GTCCGCAGAACGTCATCATCGCGAAGATCTTCCTCCTCCTGATGCGGCGCCAAATTGCAGCAAAGTAAAACAGCGTTTAGCATTGTGCT
TGTAATGATCTTGGTCAGGCAATCCGAGGGCCTGGCGTCCACAAATCTACAGAAGAACCCAAGTCTGGAGATTCAAAGAGAGGTCCCCG
AGATTCTGGGCGACTTAAGAGAAGAACAGCGCAACTCGTTCGCTGAGTTAGCGCCTTTGCAATGGGTGTAATAATTTATGTATTTTGAT
TAGGCCGAAGTTTCTGTGACTGTGGCTCGGCTTTTGCCTTTTAATTGATAAGTGTGTTTGAGATACTTTTTAATGAAGCTAATGAAACA
TGGTGAGCTTTCAGATATGTCTATATCTTTCAAGGGTTTAGGCACTCCGGTAATTTAATAAACATTTAAATCTTTAAATGAA
```

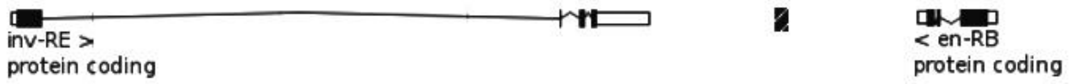


**Human EN2 superconserved**

```
Location: Chr7 - 155471753 to 155472126 (+)
TCAAACTTGAATATGCAATGAGACGCGGGAGCTTTGTGGCGGAAATATAATTAAACTGGGGAAAGATGTAAAAGCTGAAGAATGGGATG
ACATCAGGCTGATTTCACACTCGGCAGTCGGATCGCTGGGCCCAAGCCGCGCTCTTGCCACGCAAGGCAGATCAAAGTGCCCTGCCACC
GCTAAAAAGGCAAAAGGGGACTTAAGTATGCTAATCCCCAGGACAAATATATTTTAATCTTGTTAGAATACAAGTTAATGCTGCAGCTC
AGTGGCTGAACTTGGTCAGACTGTAGAGATCCATTTTTATTTTAGCTATGAATGAGCTAGACCTCTTGGTTTATTCATCGGTAATAAAA
GTAAATTTACAAAACGAA
```

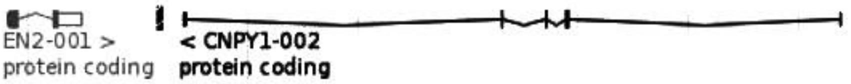


**Mouse En2 superconserved**

```
Location: GRCm38:5:28178990:28179962:1
TCAAACTTGCATATGCAACGAGGTGCTGGAGCTTTGTGGCGGAAATATAATTAAACTGGTGAGAGATGTAAAAGATGAAGAATGGGATG
ACATCAGACTGATTTCACACTTGGCAGTCCGATAGCTGGGCCCAAGCCGGGCTTGCCACGCAAGGCAGATCAAAGTGCTCTGCCACCAC
TAAAAAGGCAAAGGGAGACTTAAGTATGCTAATCCCCAGGACAAATATATTTTAATCTTGTTAGAATACAAGTTAATGCTGCAGTGCAG
TGGCTGAACTTGGTCAGGCAGGAGAGACCCATTTTTATCTTAGCTATGAATGAGGTGAAGCTCTTGGTTTTATTCATCAGTAATAAAAG
TACATTTATAAAACGAA
```

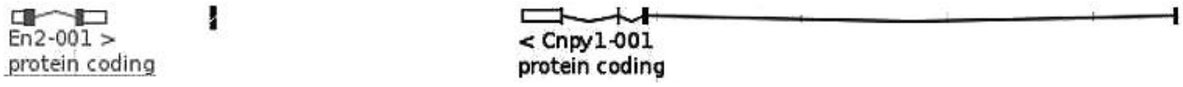


**Chicken En**

```
Location: Gallus_gallus-5.0:2:7916330:7917255:1
TCAAACTTGAATATGCAATGAAGCTCCAGAGCTTTGTGGCGGAAATATAATTAAACTGGTGAAAGATGTAAAAGATGAAGAATGGGGAT
GACATCAGGCCGATTTCACACTTGGCACTCGGATGGCTGGGCCCAAGCCGTGCTCTTGCCACGCAACACAGATCAAAGTGCACTGCCAC
TGCTAAAAAGGCAAAAGGGGACTTAAGTATGCTAATCCCCAGGACAAATATATTTTAATCTTGTTAGAATACAAGTTAATGCTGCAGCT
CAGTGACTGAACTTTGTCAGAGTGTAGAGATCCACTTTTACTTTAGCTATGAATGAGGT
```

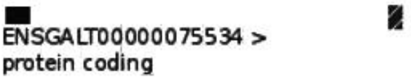


**Zebrafish eng2b**

```
Location: GRCz10:2:30000078:30000793:1
TCAAACTTGAATATGCAATGAAGAGCTGGGGCTTTGTGGCGGAAATATAATTAAGGCAGTGAAAGATGTAGAGGGTGAAGAATGGGGAT
GACATCAGGCCAATTTCACACTTGGCA
Location: GRCz10:2:30000221:30000844:1
CTTGCCACGCAACACAGATCAAA
Location: GRCz10:2:30000277:30000935:1
ATTTAAGTATGCTAATCTCCAGGACAAATATATTTTAATCTTGTTAGAATACAAGTTAA
```

Scaffold:

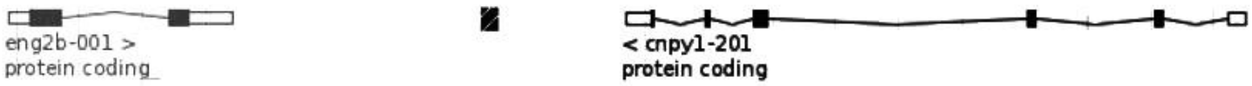


**Fugu eng2b**
Location: FUGU4:scaffold_142:748369:749065:1
GAGGTGCCCGGGCTTTGTGGCGGAAACATAATTAAAGTGGTGAAAGATGTAAAAGGTGAAGAATGGGGATGACATCAGGCCAATTTCAC
ACTTGGCA
Location: FUGU4:scaffold_142:748570:749245:1
TATGCTAATCCCCAGGACGAATATATTTTAATCTTGTTAGAATACAAGTTAATGCCGTGGCTCAGTGGCTGAACTT
Scaffold:

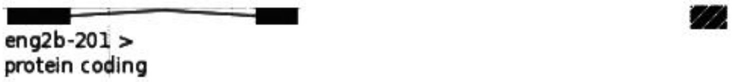


**>Human|chr7:155264047-155265809 | element 1418 | positive | hindbrain (rhombencephalon)[4/5] | midbrain (mesencephalon)[4/5]**

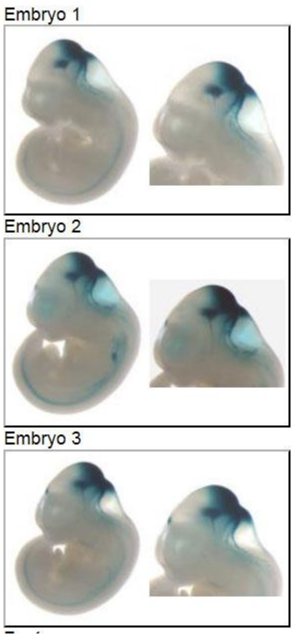


```
CAGTGACTCATCACTCGCAGAgagaaaaacatcttttatatgtagagacgggaacgcgccctgtgg
attcagagccccaaggaaacaaaaacagcacgtcagaggaagaatgtcatacaaacccgctggagt
cacaaggcaaaacagcccccatacctaaacgcccggcgtgacaatggcctttgtgcctggcgccgt
cgggggagctgggcgcctagcgggtggggcaggcggaagttggacacatttccacgcaggtgagg
ccatccatggggtgatgcgggtgttggggctcgctgctttcgctccatgaagatttgcagcccccc
ccagccaacccaccccatcccttccccccacaagggtatttcatccacactgtttttaaatgctt
cctagtcaaacttgaatatgcaatgagacgcgggagctttgtggcggaaatataattaaactgggg
aaagatgtaaaagctgaagaatgggatgacatcaggctgatttcacactcggcagtcggatcgctg
ggcccaagccgcgctcttgccacgcaaggcagatcaaagtgccctgccaccgctaaaaggcaaaa
ggggacttaagtatgctaatccccaggacaaatatattttaatcttgttagaatacaagttaatgc
tgcagctcagtggctgaacttggtcagactgtagagatccatttttattttagctatgaatgagct
agacctcttggtttattcatcggtaataaaagtaaatttacaaaacgaacccctctttgggctgtg
tcacccccaccttgagctctccttactggggactggggaattccagaggcttccaagatgggagca
tccctaacaccatccatccttctccgacctgtccccttctgtagagcatcctgatggccgaggaag
gaactctgctcacgcagcagagaagaaacagggaaactcaaaaattaattaaaagattctccaaag
caatctgtttccctttaggcttgtggtttgacaaccaaaccaattcctttctgtcagccctctcac
tttaatttttgcaatgtcaattttcattactattaattatttgccaagctccaactgtacatttgg
cattgctctgatgagcaagaagctctcaggttcctccctctttaagtgtcgattcctggaagatat
gaatatgcaagtttctgcgtgtgtgtacatgtaaaatattgggcacacatagaagaagaaatgtca
gcctgtagagctccccttgaaataaaaccactgtggtaaaatattaagcagaaccacggatatatc
tgcacagaggaatgtgtgtgtgtgtgtgtgtgtgtgtgagagagagagagagagagagagagagag
agagagagagcgagccagccagcgagccagcacttaactgtaaggttacagaaggttaaacggccc
tgcagagcctgcctgctaaatattgctgatggcagccgcgtctaaaaggcaatttctgtctttctg
ctctagcgtgtgttagaattccctctgcctcaggcagaaaacctgactcccactaaagtgtgctgc
acagccctgcattccaaagcctgtctgtgcccttccggctccctggcctcagcagggggggcgggtgcaggaggctatgcgcgcacaca
cagtcccccaggcaggccaggcaactggcagcccgaggggaggggcagctggcagcagtatccccctcccCCACAGAGAGCTTGGAAGG
T
```

*Left, intergenic sequence of hs1418 enhancer element, superconserved sequence highlighted in red. Right, three examples of hs1418 driven lacZ expression in mouse E11.5 brain specific to MHB (VISTA enhancer database).*

**EMBOSS MATCHER human EN2 consensus *inv/en* superconserved with hs1418**

```
# Length: 374 - # Identity: 374/374 (100.0%) - # Similarity: 374/374 (100.0%)
# Gaps: 0/374 ( 0.0%) - # Score: 1870

EMBOSS_001           1 TCAAACTTGAATATGCAATGAGACGCGGGAGCTTTGTGGCGGAAATATAA      50
                       ||||||||||||||||||||||||||||||||||||||||||||||||||
EMBOSS_001         402 TCAAACTTGAATATGCAATGAGACGCGGGAGCTTTGTGGCGGAAATATAA     451

EMBOSS_001          51 TTAAACTGGGGAAAGATGTAAAAGCTGAAGAATGGGATGACATCAGGCTG     100
                       ||||||||||||||||||||||||||||||||||||||||||||||||||
EMBOSS_001         452 TTAAACTGGGGAAAGATGTAAAAGCTGAAGAATGGGATGACATCAGGCTG     501

EMBOSS_001         101 ATTTCACACTCGGCAGTCGGATCGCTGGGCCCAAGCCGCGCTCTTGCCAC     150
                       ||||||||||||||||||||||||||||||||||||||||||||||||||
EMBOSS_001         502 ATTTCACACTCGGCAGTCGGATCGCTGGGCCCAAGCCGCGCTCTTGCCAC     551
```

```
EMBOSS_001        151 GCAAGGCAGATCAAAGTGCCCTGCCACCGCTAAAAAGGCAAAAGGGGACT    200
                      ||||||||||||||||||||||||||||||||||||||||||||||||||
EMBOSS_001        552 GCAAGGCAGATCAAAGTGCCCTGCCACCGCTAAAAAGGCAAAAGGGGACT    601

EMBOSS_001        201 TAAGTATGCTAATCCCCAGGACAAATATATTTTAATCTTGTTAGAATACA    250
                      ||||||||||||||||||||||||||||||||||||||||||||||||||
EMBOSS_001        602 TAAGTATGCTAATCCCCAGGACAAATATATTTTAATCTTGTTAGAATACA    651

EMBOSS_001        251 AGTTAATGCTGCAGCTCAGTGGCTGAACTTGGTCAGACTGTAGAGATCCA    300
                      ||||||||||||||||||||||||||||||||||||||||||||||||||
EMBOSS_001        652 AGTTAATGCTGCAGCTCAGTGGCTGAACTTGGTCAGACTGTAGAGATCCA    701

EMBOSS_001        301 TTTTTATTTTAGCTATGAATGAGCTAGACCTCTTGGTTTATTCATCGGTA    350
                      ||||||||||||||||||||||||||||||||||||||||||||||||||
EMBOSS_001        702 TTTTTATTTTAGCTATGAATGAGCTAGACCTCTTGGTTTATTCATCGGTA    751

EMBOSS_001        351 ATAAAAGTAAATTTACAAAACGAA    374
                      ||||||||||||||||||||||||
EMBOSS_001        752 ATAAAAGTAAATTTACAAAACGAA    775
```

**MLAGAN *inv/en-EN2* superconserved CNS**

```
sequence1(+)    000006950  ACT------GCAGACTGCAGACTGAAAACTGAAGACTGCAACCAGGGACC  000006993
sequence2(+)    000007120  TTTTAAAAGAAATGCTTCCTGGTCAAACTTGCATA-TGCAACGAGG----  000007164
sequence3(+)    000007093  GTTT--TTTAAATGCTTCCTAGTCAAACTTGAATA-TGCAATGAGA----  000007135

sequence1(+)    000006994  ACACACGACGATCGAGGCTGTGCCACAAGCTGCTCTGGGCAACG------  000007037
sequence2(+)    000007165  -TGCTGGAGCTTTGTGGCGGAAATATAA-TTAAACTGGTGAGAGATGTAA  000007212
sequence3(+)    000007136  -CGCGGGAGCTTTGTGGCGGAAATATAA-TTAAACTGGGGAAAGATGTAA  000007183

sequence1(+)    000007038  --------GAAGGCGGCAACATCGACAGTAGTTTTGCGGCTCGTTTGCTG  000007079
sequence2(+)    000007213  AAGATGAAGAATGGGATGACATCAG---------------------ACTG  000007241
sequence3(+)    000007184  AAGCTGAAGAATGGGATGACATCAG---------------------GCTG  000007212

sequence1(+)    000007080  CGTTGACTCTTCGCCGAGTCCCGTGATCATGGACATGATCATCATGACGT  000007129
sequence2(+)    000007242  ATTTCACACTTGGC--AGTCC-----------------------------  000007260
sequence3(+)    000007213  ATTTCACACTCGGC--AGTCG-----------------------------  000007231

sequence1(+)    000007130  ACGATGATCGTGGGGTCCGCAGAACGTCATCATCGCGAAGATCTTCCTCC  000007179
sequence2(+)    000007261  -----GATAGCTGGGCCCAAGCCGGG----------------CTT-----  000007284
sequence3(+)    000007232  -----GATCGCTGGGCCCAAGCCGCGCT--------------CTT-----  000007257

sequence1(+)    000007180  TCCTGATGCGGCGCCAAATTGCAGCAAAGTAAAACAGCGTTTAGCATTGT  000007229
sequence2(+)    000007285  -----------------------GCCACGCAAGGCAG-------------  000007298
sequence3(+)    000007258  -----------------------GCCACGCAAGGCAG-------------  000007271

sequence1(+)    000007230  GCTTGTAATGATCTTGGTCAGGCAATCCGAGGGCCTGGCGTCCAC---AA  000007276
sequence2(+)    000007299  ------------------------ATCAAAGTGCTCTGCCACCACTAAAA  000007324
sequence3(+)    000007272  ------------------------ATCAAAGTGCCCTGCCACCGCTAAAA  000007297

sequence1(+)    000007277  ATCTACAGAAGAACCCAAGTCTGGAGATTCAAAGAGAGGTCCCCGAGATT  000007326
sequence2(+)    000007325  AGGCAAAGGGAGACTTAAGTATGCTAATC---------------------  000007353
sequence3(+)    000007298  AGGCAAAAGGGGACTTAAGTATGCTAATC---------------------  000007326

sequence1(+)    000007327  CTGGGCGACTTAAGAGAAGAACAGCGCAA-CTCGTTCG---CTGAGTTAG  000007372
sequence2(+)    000007354  --------CCCAGGACAAATATATTTTAATCTTGTTAGAATACAAGTTAA  000007395
sequence3(+)    000007327  --------CCCAGGACAAATATATTTTAATCTTGTTAGAATACAAGTTAA  000007368

sequence1(+)    000007373  CGC--CTTTGCAATGGGTGTAATAATTTATGTATTTTGATTAGGCCGAAG  000007420
sequence2(+)    000007396  TGCTGCAGTGCAGTGGCTG-------------AACTTGGTCAGGCAGGA-  000007431
sequence3(+)    000007369  TGCTGCAGCTCAGTGGCTG-------------AACTTGGTCAGACTGTA-  000007404

sequence1(+)    000007421  TTTCTGTGACTGTGGCTCGGCTTTTGCCTTTTAATTGATAAGTGTGTTTG  000007470
sequence2(+)    000007432  -------------------------------------------------G  000007432
sequence3(+)    000007405  -------------------------------------------------G  000007405

sequence1(+)    000007471  AGATAC-TTTTTAATGAAGCTAATGAAACATGGTGAGCTTTCAGATATGT  000007519
sequence2(+)    000007433  AGACCCATTTTTATCTTAGCT-ATGAATGAGGTGAAGCTCTTGGTTTTAT  000007481
sequence3(+)    000007406  AGATCCATTTTTATTTAGCT-ATGAATGAGCTAGACCTCTTGG-TTTAT  000007453
```

```
sequence1(+)     000007520  CTATATCTTTCAAGGGTTTAGGCACTCCGGTAATTTAATAAACATTTAAA  000007569
sequence2(+)     000007482  TCAT-----------------------CAGTAA-TAAAAGTACATTTATA  000007507
sequence3(+)     000007454  TCAT-----------------------CGGTAA-TAAAAGTAAATTTACA  000007479

sequence1(+)     000007570  TCTTTAAATGAATTTACTATGGCGATACGT------TGGCTTGACATCTT  000007613
sequence2(+)     000007508  -----AAACGAACCTCTCTTTGGACTGTGTCACCCCCACCTTAAGCTTCC  000007552
sequence3(+)     000007480  -----AAACGAACCCCTCTTTGGGCTGTGTCACCCCCACCTTGAGCTCTC  000007524
```

***D. melanogaster invected and engrailed* gene loci and homology to other insects**

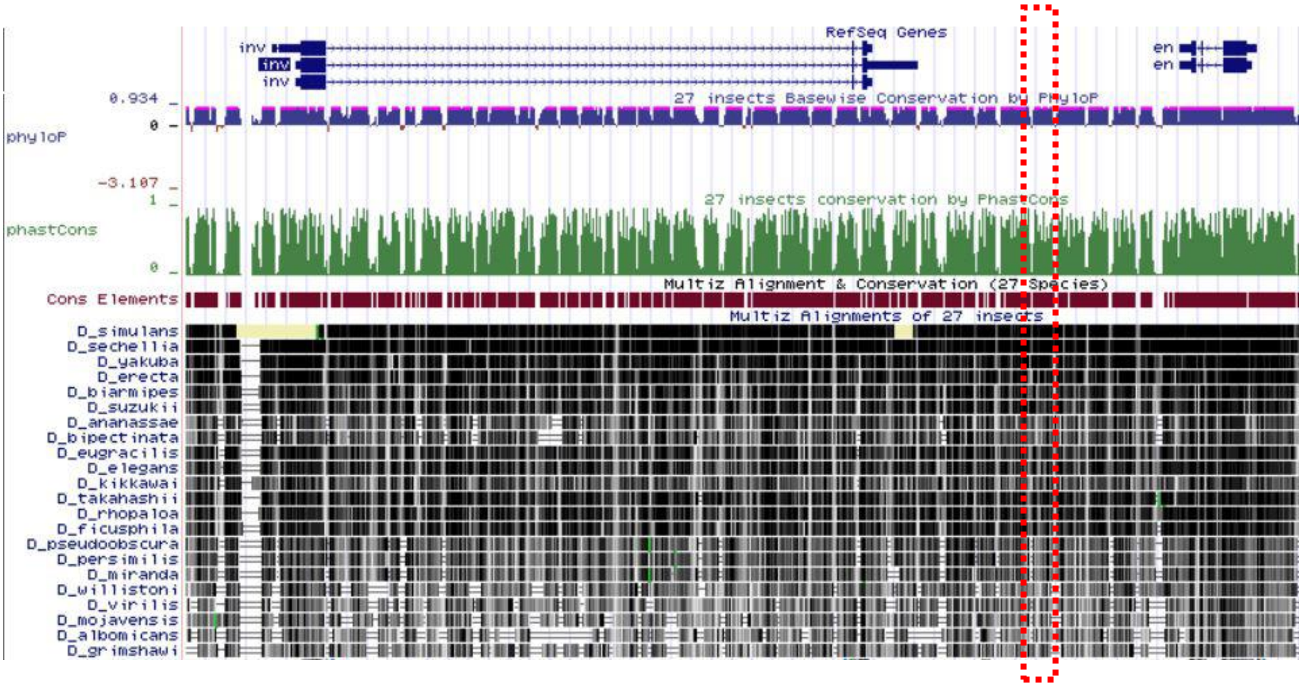


Image capture of UCSC Genome Browser on *D. melanogaster* Aug. 2014 (BDGP Release 6 + ISO1 MT/dm6) assembly, shown are the *invected* and *engrailed* gene loci. RefSeq Genes shows exon (blue bars) and intron structure; in red is shown Ensembl Gene annotation used for genome browsing. Green bar diagram indicates conservation among *Drosophilidae* species which are listed further below (black bars indicate sequence homology). Conserved elements are shown in dark red. Dashed red box indicates topology and extension of *inv/EN2* superconserved CRE sequence determined in *Drosophila melanogaster*.

**Examples of *inv/en-EN2* superconserved sequence found in other *Drosophilidae***

***D pseudoobscura* scaffold**

>chromosome: GCA_000001765.2:3:4589216:4590358:1
**GGGCAACGGAAGGCGGCAACATCGACAGTAGTTTTGCGGCTCGTTTGCTGCGTTGACTCCTCGC**TGATCATCATCATCGTGCTCATCAT
CATCATCCACAGCCACATCCACAGCCACAGCCACATCCACATCAGGCTCCTCGACGGGTCTGGTCTTGTGTCTGGTGGTGC**TGCGGCGC**
**CAAATTGCAGCAAAGTAAAACAGCGTTTAGCATTGTACTTGTAATGATCTTGGTCAGGCAATCCG**CAGACTGCAATCTTCTCGAATGGC
ACATCCAGCGAAACGGAAGAGAGTGAGCCCCAAAGATACAGCACAGAGCTTAAGAGAATGCCAGAGAGGAGGAGACAGAGAGAGAGAGA
GAT**ACTCGTTCCCTGAGTTAGCGCCTT**GGAAA**GGGTGTAATAATTTATGTATTTTGATTAGGCCG**CAAGTCGCAGACTGAAACTGAGAC
GAAGACAGAGACAGAGTCAGAGTCGGAGGCTGCATCTTCAAGTATCTTTGAGATGCACAAATTGGA**TTTAATTGATAAATGCATTTGAG**
**ATATATTTT**

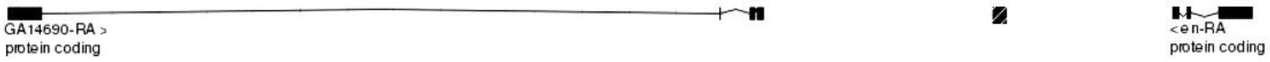


***D simulans* scaffold**

>chromosome:GCA_000259055.1:2R:5926335:5927548:1
**TGAAGACTGAAGACTGCAACCAGGGACCACACACGACGATCGAGGCTGTGCCACAAGCTGCTCTGGGCAACGGAAGGCGGCAACATCGA**
**CAGTAGTTTTGCGGCTCGTTTGCTGCGTTGACTCTTCGCCGAGTCCCGTGATCATGGACATGATCATCATGACGAACGATGATCGTGGG**
**GTCCGCAGAACGTCATCATCGCGAAGATCTTCCTCCTCCTGATGCGGCGCCAAATTGCAGCAAAGTAAAACAGCGTTTAGCATTGTGCT**
**TGTAATGATCTTGGTCAGGCAATCCGAGGGCCTGGCGTCCACAAATCTACAGAAGAACCAGAACCCAGGTCTGGAGATTCAAAGAGAGG**
**TCCCCGAGATTCTTAAGAGAAGACCAGCGCAACTCGTTCGCTGAGTTAGCGCCTTTGCAAAGGGTGTAATAATTTATGTATTTTGATTA**

**GGCCGAAGTTTCTGTGACTGTGGCTCGGCTTTAGCCTTTTAATTGATAAGTGCGTTTGAGATACTTTTTAATGAAGCTAATGAAGCATG**
**GTGAGCTTTCAGATATGTCTCTATCTTTCAAGGGCTTAGGCATTGGGGTAGTTTAATACACATTTAAATCTTTGAATGAA**

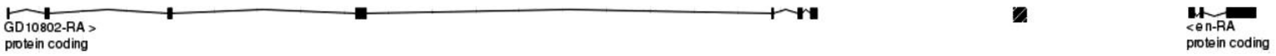


***D yakuba* scaffold**

>chromosome:GCA_000005975.1:2R:9558232:9558991:-1
**TAATATGTATGCGATTCGGCGGCAGCCGCAAAAGCATCTGTGCGGAAACTAAGAGCGAGGATCGCCGGGCGAAGAGAGGCTCCGTTTTG**
**GCAATGGTC**GATGGTCGATGCGAGTCCCCGAAAAATGCAC**TGCACTTAAAATTATTGACACTTTG**GCGATC**GATGCGGCTACGTGAGGC**
**GGCTACCGTGATGCGGCTAAGAGAGCGGTTAAGAGAGCGAAAAGGGCGGAAGAATCCAGCCGAAAGACAGGCGGGCAGACAGAAGACTG**
**CAAACTGAAGACTGC**AGACTGAAGACTGCAAAC**TGAAGACTGAAGACTGCAACCTGGGACCACACACGACGATCGAGGCTGTGCCACAA**
**GCTCCTCTGGGCAACGGAAGGCGGCAACATCGACAGTAGTTTTGCGGCTCGTTTGCTGCGTTGACTCTTCGCCAAGTCGCTTGATCATG**
**GACATGATCATCATG**

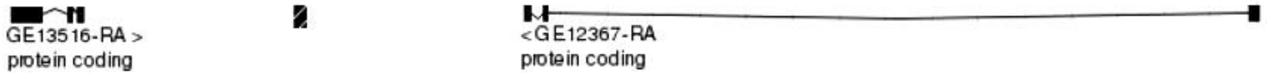


***D willistoni* scaffold**

>supercontig:GCA_000005925.1:scf2_1100000004558:2013519:2014160:-1
**AACGATCATCATGATGACGGT**TGTTGGGACG**GCGGCGCCAAATTGCAGTAAAGTAAAACAGCGTTTAGCATTG**CAGACTCGTCGTAATG
ATCTTGGTCAGGCAATGTGGCAATGTGCGTTAAATGAAGAGAAAATCTTCCCCAGAGAGAT**TTAAGAGAGTCTTCAAAGTGCTAAA**

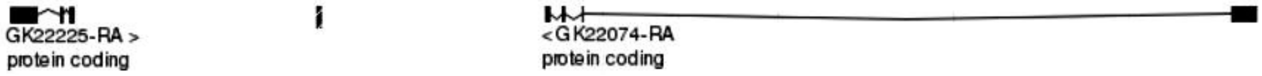

### SUPPLEMENTAL DATA SET S3

#### *dachshund/DACH1* superconserved sequence

**R65A11 - dac**

```
>BDGP6:2L:16481227:16484138:1
CGAAAATATAGCCCTTTTTCACGCTCTATTTATAGCATTCACATTCTTTCTTTTTTTTTGCACTTTTTAGCTGGCATATCCTTTCGACT
TCCGCCATTCGAGGCTCGCCCAATTTCCGTTTCGAGTTTAATTAATTTAATAAACAAATTCTTTTCGCTCTAAAAACTCTCAAGTGTAT
CGATACGATGCGTTTCTTTTTTTCCTTCGTTAAATAAATAATAACCAAAAAAAAAAAAAACCAAAAAGTAGGAGGAGAAAAGTTATTGC
CATAGTTTTTTTATTATACTTGTGTGTTTACCTTTCTGGTGGCTTGATCGATAGGCATCTGCAATTAAAAAGAGAAGAAGAAGAGACAA
GTGAGGCAAAATTGTTAAACGTTTTGTGTAAGCTTTAATACGAAAAACAAGTACTGCAACATAACGGAAGGAAACACGGCTTAAATTCG
GGGCACAAATGCTGAAAGGGAAGTTTTTCATTGACGGGTTCGTTCTGACGGACTTGCATTTTGGCGGGCAAGCGGGTGTGAAAATGCAC
ACGCCCCGAGAACCCCCCTTTCCACCCCCCCTGGACCCCTTTATCCAGCCCACTGGCCAAAAACAATTTGTAATTATCCACAGAGAGCG
CTGCCTTCAGCGGTTTCGCATTTCCCCTTTCGCTCGCTCTCCCAACTTGTTTCAATTTAGCGCAAAACTTTTTCAACCTAATAATAGGT
TTAACCGCATTTTTAACCGTTCCTCATGTTCGGTCCGGTTCGGTTTTCAAAACCGGGAATCGTACTTAGACTGGGTCTCCTTATTTCTG
TTCTGGCTCTCTGTACAACTTTTCATTGAGAAAAATGTAACTAGTTTTTCATAGCAACGGAATACAATTTAATCCAATAATCCAATAGT
TTAATCCAATACAAATGATATTACTACCATTTCTATTTTCGTTAATTTCGATTTGACTTATTTGGCTGGATTTACTTTTCAAAATATAT
GTTATCAATAAGACACAAACCTTACTTTTCTAGCTATTAACATAGTTTAAAAAAAAAAAAAAAACTAATAAAAATTACGTGAATCTAAAT
TTTTAAACCCGATATCCAAGAAGATCTCAATTTTTGCCTGTGTACTCAGTTCTCTGAACAAAGCGCATGTGCACTTTGGAGCACACTCC
ATACATGTGGCTCAGCCCTTTTCCATAATTAACTAGATGGTTTTCCATCGACTTCATTGTGGTCAGCGGCCAGTTCAACCCGTTCTTCA
CTGCAACCGAGAACTGTAAACACAAAAACCCCAGGACTCTACATTGGCTTAAAAAAATAAGAACAGAACAGAACCAAAACCAAAAAAAG
AAGGGATATTGAAATACAAGGTTGTAATCGTTTCGACTGTTGATGTCTCAATGCATGGGCAGTTCAGTTAGTAAATGTTTTTCAAAATC
TTTCAGGCAGGAGATGTTTAAATATCCATGAAATATTTGGATCTCCTGGGGATCAATCGGAATATTAGCCTTTAATTGTGTTGATCTTT
TAAGCCTTTTTGTATCTAATCTAAGCCATTCGATCTAATCACAATTTATAAATATCTGCATATTTCTGTATAAGTCTGCATCATTTGAC
GTAACTCTTTAAGTCTTTTGGCTTAAGTTGCAACTATAAGGAAGTATTTATTTTAGAGACACAAATATTTCAGTCGCCTTCATTTGAAC
AAATCGGCGAAAATTGGCTAGCTCGCCAAACTTTCTGTAACCAAGGACAATGGTTTTATTTTAAACCATTAAAAACTTTAGACCCACTA
GCTCCTAGATCCCCCTCAAAAGATTTAAAAAAAAAAAACACGATACCCATTTCTACTGAACTTCGTTTTTGCTTGTCGTTTTTTCCACT
CGAACGGAAATGAGCTGACAGCGCACCGCACACGTCGATTGCAGAAAAACATCGGATAAAACAGGAGGAAAAGTTGTGCAAGGTGGAAA
ACTGTTTTACCAACTATTGTTAGAGGCGTTCAAAAGAATTACGCAGCTTTCGGTTAGTTAGCAAGGGGTCACCGGGGAGCGTTACGTTT
GCATTGCGTATTTCCGCTAAATGTCATCGGAAAAGGCAAACGGCGAAATGCGAAACGAAAGTTTTTTGATTGCCCGTGTTAATCGATAT
CGATGCACAAACTATTTGCATTGCAACCGTTGCAAGAATATGCAAGAAGTTGGGGGCGGCCGCGGCAGGGGGTGGAAGTTGAGTGCGTA
AGTTGGCTAAAGCGGAAACAGGAAATGAGAAAATTTTGCAGAGCAAACCCCGAACTGGAAATGCAACTAACTGGGCACATGCACTTTGC
GAAATCATTGGATAGCGTTAAGAAATTTATTTTAAAATTGTAACTAACATTTAATCGTATTCAAAAGCAATTAAATCCCAATCCAATTC
TTATATAAAATCCTTACAAGATTATTCTATTTACTGTAAATCTAAGCAAAAACTCCCTTTGCAAAATATTCGCCTGCACAGCACAGATC
AGTGAAATAATCAAATGAAGTCTTGAAATAACGAAAAACCCCCAATTGCGTGTGGAACTGCCCCCAATGCTTTTGCTTCGGTTTCGTAC
CTGGCCGTGGTGCAGTCCCTGTAGAGGATGTCGAAGTCCTTGCAGCAGAGCAACTTGCAGCGATTGACGCCCGGCTGGATGGCGCCCAA
GCCGCGTAATATGCGCACCTGTTCCACATTGCAGACGAGCGGTACGATGTCAAGGCGCTTTAGTTTGGTATAAACCGTATGCAGACCGC
CGACCAAGTGCTTCAGGAAGAGCTCGAAGGCCTGCGGCAGGCAGAGCATCGTTTCGTTGCTAATTATGAATGCGGCGACCTTCTGACCC
CGGTACTCCACCAGCTTGCACTCATTGGCACTGGGATCCGAGGTGGAGATCGGCGGCGGTGAGT
```

*Yellow highlighted sequence depicts dac superconserved sequence*

**EMBOSS MATCHER R65411 - dac superconserved**

```
# Length: 247 - # Identity:     247/247 (100.0%) - # Similarity:   247/247 (100.0%)
# Gaps:             0/247 ( 0.0%) - # Score: 1235
R65A11dac         79 TCCTTTCGACTTCCGCCATTCGAGGCTCGCCCAATTTCCGTTTCGAGTTT    128
                     ||||||||||||||||||||||||||||||||||||||||||||||||||
Dacyellow          1 TCCTTTCGACTTCCGCCATTCGAGGCTCGCCCAATTTCCGTTTCGAGTTT     50

R65A11dac        129 AATTAATTTAATAAACAAATTCTTTTCGCTCTAAAAACTCTCAAGTGTAT    178
                     ||||||||||||||||||||||||||||||||||||||||||||||||||
Dacyellow         51 AATTAATTTAATAAACAAATTCTTTTCGCTCTAAAAACTCTCAAGTGTAT    100

R65A11dac        179 CGATACGATGCGTTTCTTTTTTTCCTTCGTTAAATAAATAATAACCAAAA    228
                     ||||||||||||||||||||||||||||||||||||||||||||||||||
Dacyellow        101 CGATACGATGCGTTTCTTTTTTTCCTTCGTTAAATAAATAATAACCAAAA    150

R65A11dac        229 AAAAAAAAAACCAAAAAGTAGGAGGAGAAAAGTTATTGCCATAGTTTTTT    278
                     ||||||||||||||||||||||||||||||||||||||||||||||||||
Dacyellow        151 AAAAAAAAAACCAAAAAGTAGGAGGAGAAAAGTTATTGCCATAGTTTTTT    200

R65A11dac        279 TATTATACTTGTGTGTTTACCTTTCTGGTGGCTTGATCGATAGGCAT    325
                     |||||||||||||||||||||||||||||||||||||||||||||||
Dacyellow        201 TATTATACTTGTGTGTTTACCTTTCTGGTGGCTTGATCGATAGGCAT    247
```

**hs137 DACH1 intragenic enhancer**

```
GCAATTTTGAAAAAGAAAACAATGGTTAGGAAGACTTTATTTTATTCTAATTATATTTGATACTTTAAATGTTTATCTATCTAATATAT
CTCTTATTAAACAATTCTTTAGCTCTGGTATATTTTTTAGTAAAGTTTTAATAACTTTCTCAGAATTAAAAATAGCATATGATATCAGA
```

GACCAACAATAAAAGTAATCGAATTTAGAGAAATGTTTAGAAAATAAATAGATCTATACATATTTTGTTAAAGACCAATCAATTGATCT
GCTGTTTTCTATGTAAAATGAAACAATGCTTATTATTAAAATATAGGCTGAATTCTTTGCCTGATTTTCCGCTTTTGTAAAATGCATTG
CTTGAGAAACAAGTCTTCAAAAAACAAGGCCTAAAAAAATCTACTACTCAGGGTGCCTTTGAGGATATTTAATTAAAATCTAATCCTGC
ATTCATTAAGGCTCACATAAATTAAGCTGTCATTCATAAGATTTATGGATTCTCATTTGCATATTGCATACAATTCATCAATTACTCAA
GTATGAAAGGAGCACATTTCCCTTGGAGCTGCCTGCTACCCTGCCAACATTTGAAATGAGGGAAAGAGCAAGACTGTCAGGCATTCACA
CAAACTTTCTTCCAAATGTCTGCTCCTTGATTAATCTAATTTTCTAGATCTTCCCTACAAGATACACCAACAGCCCTGGTGCACATTTC
TATTATTTCGCCTGTCTTCATCACCAAAATCTTCCAGTCTTTGAATACTTCTTCATGTCTGAGTGATTATTCTTTATTTGCTAATAATA
TGGCATTGATGATATTTCCCTATTCAAATGTGTTTGATATGTACAAAGATGTAGGTTATAGGTCTGTTGATTTTTAATGTTTATTATTT
GCTATCAACAATCAGCCAATAAAGAGCTGAAAATGAGGACATGGAAAAGGAAGAGAATGTTGCTGTGGAATGTAGTTGGCACAAAAGCC
TAGTTTCAGAGACTGTTAGCATTTCAGAAGAACCTGAAACAAAGCCCTGTAGTAAATCACTGATTCTATATTTAATATAAAAAATGGAG
TTCACTCAATCAGCTGCAAGTATATGCTACTGCACTTTTTGCAGTTCTAAGCAAAGTAGTAAGTAACCCAAATTAGTTTGTTTTCACTG
TGCATAAAGCATACTGTGTTAATTCACTTGTCTTCTCTGGGGAGTAGATCTTAATTTCAGTGGCACAAGTTGGAAGTAACAGTCCTTTC
TAGACCACACTCTGGAAACCTTCCAACAGGAATGTGATCAGGTGCGAAACTTAATGCAAAACACTCCTCATGTCCAGGATTTCTACAAT
TAAAATATATGAATGTATCAGCACTCTCAAAAATCTCTGCTCCTGAAAACATGCTGTCAATTAAATGTCAATTAAATAGAAAAACGGGA
AAAACTACCTTTCAAAATCATTAGTTTTCTATGAAGTGCATATACTTCAGTATAAAAAGAAAGCATTTTATTCAATTTAAATATTTAGA
AAATTCATTTGGTAACATGAGGGATTTTATTTTACTGTATGATACTTGCCAAAACTATGATCATATCTCTTTGGAAACACCCATTTAGA
ACCCAAAATTAGAATTTCCCAAATTAAAACCTTAACATCTCAGGTTAAGTGTTAAATTACCAAAGACACCTTTTGGTTGATTTTGCTCC
CGAAGAGTCTAAAGAAATGAGTAAATGACTCTAAGGACTATACTAGCATAATCATAAATATTTTCACTTTACAAAAACTGTGCCTCTAA
AGAGAAAAAAATTGTATTCAGCAATAAAAATAATTTCCATGAATTCCTCACAAATACATTTAACATAGGAGAAATAAATGACTCTATAA
ATTTAATCAGAAGGATGAAATCATTAGCATCCCCAATTTATAAAATGACTTTCAAAACTGAAAGCAAACAACAGATAAGCTCATGCTTT
TTATTCATTCAAGTGGGATGGTTCTTCATAATTTTCAGAGATCCATGCCTGAGACACTACTGAGTGTATTGTCTGTCTTTTAACTCTTG
CATTCTCTACATTTCAGTTTTCTCAGCATGAGCTGTCTA

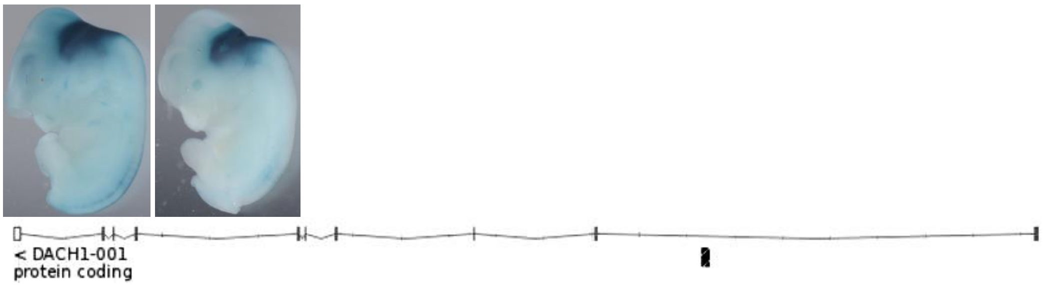


*Left, two examples of hs137 driven lacZ expression in mouse E11.5 brain specific to MHB (VISTA enhancer database); right, intragenic location of hs137 enhancer element within human DACH1 locus.*

**EMBOSS MATCHER hs137 - dac superconserved**

```
# Length: 96 - # Identity: 60/96 (62.5%) - # Similarity: 60/96 (62.5%)
# Gaps: 5/96 (5.2%) - # Score: 108

hs137DACH1int   1049 ATTTAATATAAAAAATGGAGTTCACTCAATCAGCTG-CAAGTATAT-GCT   1096
                     |||||||| ||.||||....|||.|||.|..|.||. |||||.||| |.|
Dacsupercons      56 ATTTAATA-AACAAATTCTTTTCGCTCTAAAAACTCTCAAGTGTATCGAT    104

hs137DACH1int   1097 AC--TGCACTTTTTGCAGTTCTAAGCAAAGTAGTAAGTAACCCAAA   1140
                     ||  |||..||.||....|.||..|..||.||...|.|||||.|||
Dacsupercons     105 ACGATGCGTTTCTTTTTTTCCTTCGTTAAATAAATAATAACCAAAA    150
```

**dac superconserved**
TCCTTTCGACTTCCGCCATTCGAGGCTCGCCCAATTTCCGTTTCGAGTTTAATTAATTTAATAAACAAATTCTTTTCGCTCTAAAAACT
CTCAAGTGTATCGATACGATGCGTTTCTTTTTTTCCTTCGTTAAATAAATAATAACCAAAAAAAAAAAAAAACCAAAAAGTAGGAGGAGA
AAAGTTATTGCCATAGTTTTTTTATTATACTTGTGTGTTTACCTTTCTGGTGGCTTGATCGATAGGCAT

**mDach1 superconserved**
GGTTAAAGTGAATATTTTCAGCGTGAACTTGTCTCTTAATGTCCATTAGACTGACTTTCTTGCCCTTTGTAGCACATTTGTATTCTGTG
GAGAAAAGGAAAATTGATCCCTGAGGCCACTAATGGAAACACTTTCTATACCAGTTTTCAGTTTTTCAATTGATTGAATTAGATTTTAG
AAATGTTTAGACTAACAATATATGAGTTTAACCGAAAAATAGAAGAAGAAAAACAGCCAAGAAACTGGTAGTAAATGAGCCTTTATTGC
CTCAGGCATTGGCCTAGTTAATAAACTTTCATTGAAGGTTTCTTTCAACTGTTACGGACAATGAGAGGGGGAAAAAATCTTAAATTACA
GCCATGTGTGACTTTTAAAGCAGTCATTCTTTTTTATCTCAATACAACATTCCTTTGTCTAGCATCTCCATGAAAATTATCATTTGAAA
GGTTTATATCTTAAATAAAGGGGAAAAAACCATTTTCTTCTAAATACAATTAAAACTAGTTTGAAAAGTAAGAGGCTCAGGAAACAAGA
TCTGAATAAGTAATGTATTAGCAGGTAGAACATCTTGAAAACTTACTAAGAATGCT

**hDACH1 superconserved**

GGTTAGACTGAAGATTTTGAGTGTGAACTTTTCTCTTAATGTCCATTAGACTGACTTTTCTTGGCCTTTGAAACACATTTGTATTCTGC
AGAGAAAAAGAAAAATTGATCCTTGAGGCCATTAATGGAAACACTCCCTATACCAGTTTTGAGTCTTTCAACTGATTGAATTAGATTTT
TTAAATGTTTGTACTAACAACATATGAGTTTAACCGAAAAATAGAAGAAGAAAAACAGCCAAGAAACTGGTAATAAATGAACCTTTATT
GCCTCAGGCATTGGCCTAGTTAATAAACTTTCATTGAAGGTTTCTTTCAACTGTTACGGACAATGAGAGGAAAAAAATATTAAATTACA
GCCATGTGTGACTTTTAAAGCAGTCATTCTTTTTTATCTCAATACAATATTCCTTTGTCTAGTATTTCCATGAAAATTATCATTTGAAA
GGTTTATATCTTAAATAAAGAAAAAAGTTTCTTCCAAATACAACAAAAACTAGTTTAGAGAAATAGTATGCTGAATAAATGAGATCTAA
AAATATGCTACAACCATATTCATATGTAACATCTGAAAACCTATTAAGAGTGTT

**_D. melanogaster dachshund_ gene locus and homology to other insects**

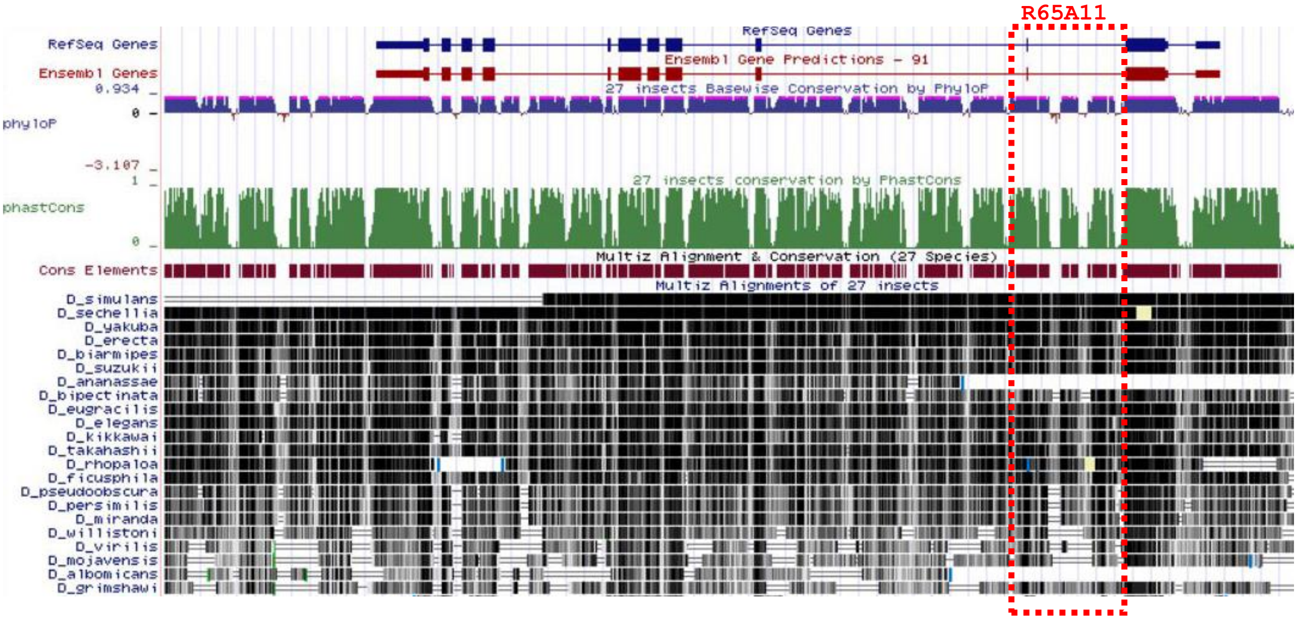


Image capture of UCSC Genome Browser on *D. melanogaster* Aug. 2014 (BDGP Release 6 + ISO1 MT/dm6) assembly, shown is the *dachshund* gene locus. RefSeq Genes shows exon (blue bars) and intron structure of dac; in red is shown Ensembl Gene annotation used for genome browsing. Green bar diagram indicates conservation among Drosophilidae species which are listed further below (black bars indicate sequence homology). Conserved elements are shown in dark red. Dashed red box indicates topology and extension of R65A11 CRE sequence determined in *Drosophila melanogaster*.

**Examples of dac superconserved sequence found in other _Drosophilidae_**

**_D melanogaster_ scaffold**

TTGTTCGCAACCAACTAACAGAGGTTCGTCTCTAACATTTTTCAAAAAAATTACATAACTTTTAAATTTGATTTCAGTTTATTTGTAAG
TGAGAAGCCTATTTTCTAACCATAAATTCTGCACGTTAAGAGTATTTCCTTTCATATCGTATCTACAAAAATCAATCCAACACACCTGT
TTCATCTACCGTTAACACCGTTAAGCCCCGCCCCATTTTCTTATCGAAAATATAGCCCTTTTTCACGCTCTATTTATAGCATTCACATT
CTTTCTTTTTTTTTGCACTTTTTAGCTGGCATA**TCCTTTCGACTTCCGCCATTCGAGGCTCGCCCAATTTCCGTTTCGAGT**TTAATTAA
TTTAATAAACAAATTCTTTTCGCTCTAAAAACTCTCAAGTGTATCGATACGATGCGTTTCTTTTTTTCCTTCGTTAAATAAATAATAAC
CAAAAAAAAAAAAA**AACCAAAAAGTAGGAGGAGAAAAGTTATTGCCATAGTTTTTTTAT**TATACTTGTGTGTTTACCTTTCTGGTGGCTT
GATCGATAGGCATCTGCAATTAAAAAGAGAAGAAGAAGAGACAAGTGAGGCAAAATTGTTAAACGTTTTGTGTAAGCTTTAATACGAAA
AACAAGTACTGCAACATAACGGAAG

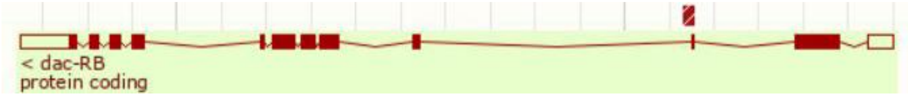


**_D sechellia_ scaffold**

>supercontig:GCA_000005215.1:scaffold_7:127389:128111:1
CATTGTTCGCAACCAACTAACAGAGGTTCGTCACTAATATTTTTCAAAAAACTTAATAATTTGCAATTTTTAATTCAGTTTATTTGTAC
TTGGGAAGCCTATTTTCTAACCATTTAATCTGCATGTTAAGAGTATTATTTCCTTTCATATCGTATTTACAAAAATCAATCCACACACC
TGTTTCATCTACCGTTAACACCGTTAAGCCCCGCCCCATTTTCTTATCGAAATAAAACCCTTTTTCACGCTCTATTTATAGCATTCACA
TTCTTTTTTTTTTGCACTTTTTAGCTGGCCATA**TCCTTTCGACTTCCGCCATTCGAGGCTCGCCCAATTTCCGTTTCGAGTTTAATTAA**
**TTTAATAAACAAATTCTTTTCGCTCTAAAAACTCTCAAGTGTATCGATACGATGCGTTTCTTTTTTTTCCTTCGTTAAATAAA**ATAATA
AC**AAAAAAACCAAAAAGTAGGAGGAGAAAAGTTATTGCCATAGTTTTTTTATTATACTTGTGTGTTTACCTTTCTGGTGGCTTGATCGA**
**TAGGCAT**CTGCAATTAAAAAGAGAAGAAGAAGAGACAAGTGAGGCAAAATTGTTAAACGTTTTGTGTAAGCTTTAATACGAAAAACAAG

TACTGCAACATAACGGAAGGAAACAAGGCTTAAATTCGGGGCACAAATGCTGAAAGGGAAGTTTTTCATTGACGGGTTCGTTCTGACGG
ACTTGCATTTT

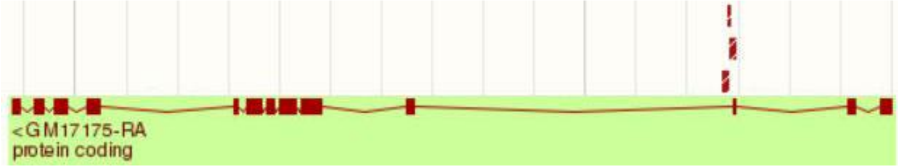


***D simulans* scaffold**
>chromosome:GCA_000259055.1:2L:16190558:16191295:1
AGGATCATTGTTCGCAACCAACTAACAGAGGTTCGTCACTAATATTTTTCCAAAAAATTAATAATTTGAAATTTTTAATTCAGTTTATT
TGTACTGGGGAAGCCTATTTTCTAACCATTTAATCTGCACGTTAAGAGTATTTCCTTTCATATCGTATTTACAAAAATCAATCCACACA
CCTGTTTCATCTACCGTTAACACCGTTAAGCCCCGCCCCATTTTCTTATCGAAATAAAACCCTTTTTCACGCTCTATTTATAGCATTCA
CATTCTTTTTCTTGCACTTTTTAGCTGGCCATA**TCCTTTCGACTTCCGCCATTCGAGGCTCGCCCAATTTCCGTTTCGAGTTTAATTAA**
**TTTAATAAACAAATTCTTTTCGCTCTAAAAACTCTCAAGTGTATCGATACGATGCGTTTCTTTTTTTCCTTCGTTAAATAAA**ATAATAA
C**AAAAAAACCAAAAAGTAGGAGGAGAAAAGTTATTGCCATAGTTTTTTTATTATACTTGTGTGTTTACCTTTCTGGTGGCTTGATCGAT**
**AGGCAT**CTGCAATTAAAAAGAGAAGAAGAAGAGACAAGTGAGGCAAAATTGTTAAACGTTTTGTGTAAGCTTTAATACGAAAAACAAGT
ACTGCAACATAACGGAAGGAAACACGGCTTAAATTCGGGGCACAAATGCTGAAAGGGAAGTTTTTCATTGACGGTTTCGTTCTGACGGA
CTTGCATTTTGGCGGGCAAGCGGGTG

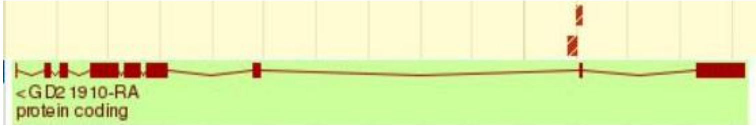


***D yakuba* scaffold**
>chromosome:GCA_000005975.1:2R:2939300:2940051:1
GAGGATCATTCTTCGCAAACAATAGAGGTATGTGTAGGTCTCTAACATTTTTCAAAATATTACTCTATTTAAAAGTGTAAATTCGGTTT
ATTTGTGTGTAACTAGCCTATCTTCTAACCATTTCTTCTGGACGTTAAGAGTATTTCCTTTCATATCGTATCTCCAAAAATCAATCCAC
ACACCTGTTTCATCTACCGTTAACACCGTTAGGCCCCGCCCCATTTTCTTATCCAAATAAAACCCTTTTTCACGCTCTATTTATAGCAT
TCACATTCTTTTTTGCACTTTTTAGCTGGCATA**TCCTTTCGACTTCCGCCATCCGAGGCTCGCCCAATTTCCGTTTCGAGTTTAATTAA**
**TTTAATAAACAAATTCTTTTCGCTCTAAAAACTCTCAAGTGTATCGATACGATGCGTTCATTTTTTTCCTTCGTTAAATAAATAATAAC**
**AAAAAAAGAACCAAAAAGTAGGAGGAGAGAAATTATTGCCATAGTTTTTTTATTATACTTGTGTGTTTACCTTTCTGGTGGCTTGATCG**
**ATAGGCAT**CTGCAATTAAAAAGAGAAGAAGAAGAGACAAGTGAGGCAAAATTGTTAAACGTTTTGTGTAAGCTTTAATGCGAGAAACAA
GTACGGCAACATAACGGAAGGAAACACGGCTTAAATTCGGGGCACAAATGCTGGAAGGGAAGTTTTTCATTGACGGCTTCCTTCTGACG
GACTTGCATTTCGGCGGGCAAGCGGGTGTGAAAATGCACA

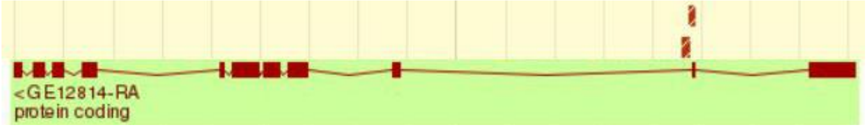


***D ananassae* scaffold**
>supercontig:GCA_000005115.1:scaffold_12916:16070598:16071275:-1
AAAATGATCATTGGGTTGGCTGCCATTTGAGAAGCCCCCAATAAAGGCAAACAAATATAATAAAAAATTATTGCAACGTAAGTTTTGAA
TACAATTGGACGGTCTCAAGAGTGTTTAGAGTGAAAAACAATAAATGGTGTTAATTTTTGTTCAGTTTTTAAAATCAAACTCAATACCT
GTTCCATCTATATACCTTTCATATCGATCCCCTTGCTCTAATAATCATGCTATATCATATTTTTTTATGTTTCATGCTCTATTTATCCT
TCCGGGCTTATCCTTTCGACTTCCGGGACCCTG**GAGGCTCGCCCAATTTCCGTTTCGAGTTTAATTAATTTAATAAACAAATTCTTTTC**
**GCTCTAAAAACTCTCAAGTGTA**CAAACTTTTTTACTGTTTTTTTGTTACCTTTAAAAAACAACGATAACCGAAAAAATTTATAAACAAA
AAATTTTCTTAGGGGGGGAAATAGGAGCGAAA**TTATTGCCATAGTTTTTTTATTATACTTGTGTGTTTACCTCTCTGGTGGCTTGATCG**
**ATAGGCAT**CTGCAATTAAAAAGAGAAGAGAAGAGACAAGTGAGGCAAATTGTTAAACGTTGTGTGTAAGCTTTTAATACAGTGAGAGAA
AAGAAAAACAGGAAACCAAAAAAAACAATATTCAGAGGGATGGTATTATGCTGCA

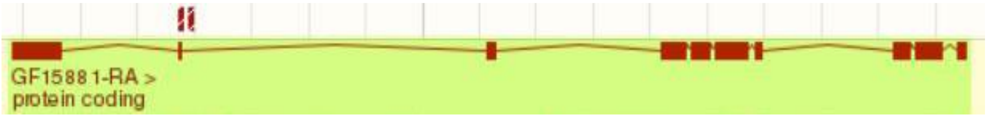


***D pseudoobscura* scaffold**
>supercontig:GCA_000001765.2:4_group3:8197417:8198097:1
AGATCTGTTCCCAGTTGTTCAAACTAAAAGTTAAGCTCAAGTTAAAAATGATCAAATTACTCCCAGACGGTATTGGATTTCCTGTCTCC
CCTTGGTTTCGACCATAGATTATCTTTCAGAACCGTTTCAATCGCTTTATCTGCGTATAAAAAATCCAATCCCCTCACCTGTTCGCTCT
ACTGTTAACACCGTTATAACCGTAATTCCCAAAAATGAAACCCAAAAATTCTTATCGGATAAACCCTTTTTTCATGCTCTATTTCTTGT
TTAACCCTCAAATATCCTTGCAACTTCCGCATT**TTCGCGGCTCGCCCAATTTCCGTTTCGAGTTTAATTAATTTAATAAACAAATTCTT**

**TTCGCTCTAAAAACTCTCAAGTGTA**GGAAAAAAACGATGCAATCGATGCGTTTCTTTCTTCCAAAAAACAAAAAAACCAAAAAATATTA
AAAAAAAAAAAGAGT**AAAAGGAGGAGGAGAGAAATTATTGCCATAGTTTTTTTATTATACTTGTGTGTTTACCTCTCTGGTGGCTTGAT
CGATAGGCAT**CTGCAATTAAAAAGAGAAGAGAAAGAGACAAGTGAGGCAAAATTTGTTAAACGTTGTGTGTGTGTAAGCTTTAAACAAC
AAAAGTAAAAAACGAGTGAAACTCTGAATAAAACAGAGGAGTTCCGTCCCCGAAAACA

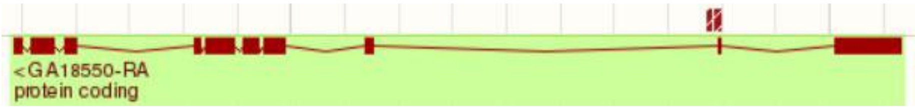


***D willistoni* scaffold**
>supercontig:GCA_000005925.1:scf2_1100000004585:4705370:4706001:1
AGAAGGCCAGCGTAACGAAATTATAACATTTTCATGCTCTATTTACGTATCCTTCGATTTCAAAAATTTGTGAATGGACTCTCCCATTT
CCGTTTTTTTTTTTTGTTTATGTAAAATATAATTTAATTTAATAAGAAAAAACTAAATTTG**TTTTCGCTCTAAAAACTCTCAAGTGT**TT
TGTGTGTATATAGAAAAAAAATGAATGTATACAATATGCGATCGATTTTTCATAACGGAAAAGATATTATTGCCGTAGTTTGTTTTTGT
TTTTTTTTTTTG**TTTTTTATTATACTTGTATGTTTTACCTCTCTGGTGGCTTGATCGATAGGCAT**CTGCAATTAAAAAGAGAAGAAGAA
GAGACAAGTGAGGCACAACATTTTTGTTAAAACATTACAACAAAAAAAAAAGGAGAACATAATGTAATAAGTAAAAGAGAAAATATTAC
ATAGGGATGTATTTTAGACGGAAGTTTCGGGGGCAAAAGCGGGTGTGAAAATGCACACGCCCCTTACCCCCGCTCCCCTTTCCCCGGCA
CATTCCTTCCCCTCCTCTGACGTCTGTCTATCTTACTGAGGTCCCTCATAATTATTATGGCCAGGGCCCAAACAGTGGGGTGCAACTTC
GACCTACCA

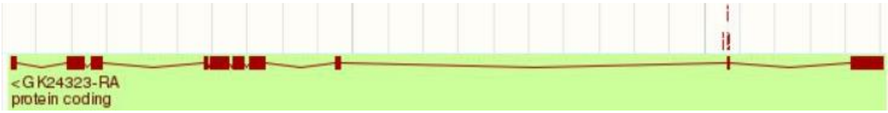


**D grimshawi scaffold**
>supercontig:GCA_000005155.1:scaffold_14978:688467:689119:1
CTCGCTTTGACCCACATCTTTTCGCATAGCTGGCCAAACGCGTTATCCTTAACCAAAAGCAACAACAAAAAAGAAAAAGAAGAGGAAAA
AAAGCCCATTAACACGCTCTATTTGTTTACGTTCCCAAAAAACTTCCGGCTGTCGTACGCTGGCTCGTCCCATTTCCGCATTAAGTAAT
TAATTTAATAACAATTTTTTATTTTTTTGTTTTTTT**TTTTCGCTCTAAAAACTCTCAAGTG**CACACAGAACCGAGTACATCGATTGTAGG
ATCGATTTCGATTTCAACAGATTGCCATAGTTT**TTTTTTTATTATACTTGTGTGTTTACCTCTCTGGCGGCTTGATCGATAGGCAT**CTG
CAATTAAAAAGAGAAGAACAAGAGACAAGAGTGAAGTATAAAAAACACATAGGTAAAAAACGTTGTATGTGTGTGTGTGGTTTGTTGTG
TGTGTGTGTGTTAAGCTTCGACTAAAACAAAAACAAAAACAGCAACAAAAAACAAAAAAAAAACACGATCAGAAAAAAGAAAACGCAAT
TTTAAAAATTTCAAAATTATGTTTCACTTATGGACGGCACGGCGGGAAAGCGGGTGTGAAAATGCACACGCCCCAACCCCTTCAGAAAA
AAAAAAGAGACGCGGGCAGCCACGATAACT

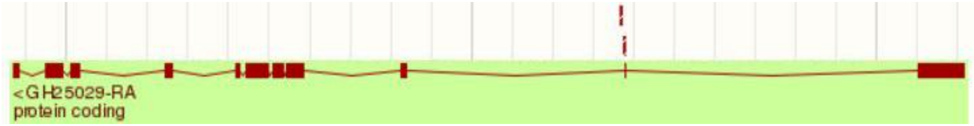
